# Gas tunnel engineering of prolyl hydroxylase reprograms hypoxia signaling in cells

**DOI:** 10.1101/2023.08.07.552357

**Authors:** Peter Windsor, Haiping Ouyang, Joseph A. G. da Costa, Anoop Rama Damodaran, Yue Chen, Ambika Bhagi-Damodaran

**Affiliations:** Department of Chemistry, University of Minnesota, Twin Cities, Minneapolis, MN, 55455, United States; Department of Biochemistry and Molecular Biology, University of Minnesota, Twin Cities, Minneapolis, MN, 55455, United States

**Keywords:** non-heme iron, cellular signaling, oxygen sensing, gas tunnels, protein design

## Abstract

Cells have evolved intricate mechanisms for recognizing and responding to changes in oxygen (O_2_) concentrations. Here, we have reprogrammed cellular hypoxia (low O_2_) signaling via gas tunnel engineering of prolyl hydroxylase 2 (PHD2), a non-heme iron dependent O_2_ sensor. Using computational modeling and protein engineering techniques, we identify a gas tunnel and critical residues therein that limit the flow of O_2_ to PHD2’s catalytic core. We show that systematic modification of these residues can open the constriction topology of PHD2’s gas tunnel. Using kinetic stopped-flow measurements with NO as a surrogate diatomic gas, we demonstrate up to 3.5-fold enhancement in its association rate to the iron center of tunnel-engineered mutants. Our most effectively designed mutant displays 9-fold enhanced catalytic efficiency (*k*_cat_/*K*_M_ = 830 ± 40 M^-1^ s^-1^) in hydroxylating a peptide mimic of hypoxia inducible transcription factor HIF-1α, as compared to WT PHD2 (*k*_cat_/*K*_M_ = 90 ± 9 M^-1^ s^-1^). Furthermore, transfection of plasmids that express designed PHD2 mutants in HEK-293T mammalian cells reveal significant reduction of HIF-1α and downstream hypoxia response transcripts under hypoxic conditions of 1% O_2_. Overall, these studies highlight activation of PHD2 as a new pathway to reprogram hypoxia responses and HIF signaling in cells.

## Introduction

Oxygen (O_2_) is a critical metabolite required for oxidative phosphorylation and subsequent energy generation. Cells utilize diverse mechanisms to recognize, respond and adapt to change in O_2_ levels in order to properly function.^[1–5]^ Humans and other mammals utilize non-heme iron-based O_2_ sensors that function by using O_2_ as a co-substrate to catalyze the hydroxylation of specific amino acid residues of partner proteins.^[6]^ A prime example of an O_2_ sensing non-heme iron sensor is mammalian prolyl hydroxylase 2 (PHD2) that utilizes O_2_ to catalyze hydroxylation of transcription factor, hypoxia-inducible factor HIF-1α (left panel, **Fig. 1A**).^[7,8]^ The O_2_ sensing capability of PHD2-like non-heme iron-based sensors is often discussed in terms of their catalytic efficiency with respect to O_2_ (*k*_cat_/*K*_M_(O_2_)), and can vary dramatically depending on the enzyme (**Table S1**).^[9–11]^ For example, PHD2 has a *k*_cat_/*K*_M_(O_2_) of approximately 90 M^-1^ s^-1^, AspH has a *k*_cat_/*K*_M_(O_2_) of approximately 550 M^-1^ s^-1^, whereas factor-inhibiting hypoxia-inducible factor (FIH) has a *k*_cat_/*K*_M_(O_2_) of 5100 M^-1^ s^-1^ (**Fig. 1A**).^[9,10]^ The structural and molecular features that govern O_2_ sensing capability of PHD2-like non-heme iron-based sensors are not fully understood.^[12]^ This gap in knowledge remains despite the prevalence of these sensors in bacteria, single-celled eukaryotes, and animals (**Fig. 1B, Fig. S1**).^[11,13,14]^

**Figure 1.**
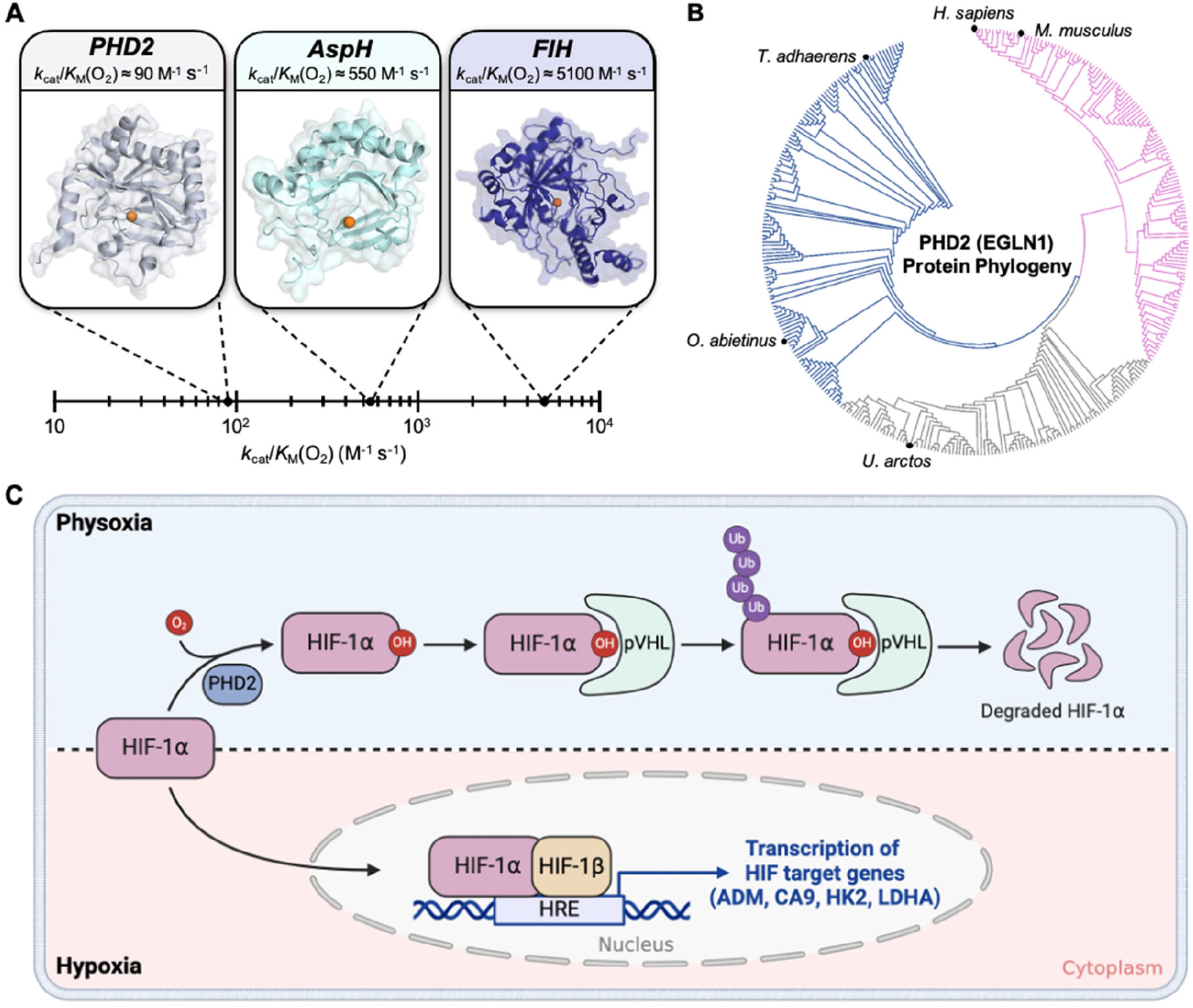
**A)** Range of catalytic efficiencies with respect to O_2_ (*k*_cat_/*K*_M_(O_2_)) for O_2_ sensing non-heme iron hydroxylases in *H. sapiens*. **B)** Protein phylogeny of PHD2 (also referred to as EGLN1) across 1844 organisms. PHD2 is well conserved throughout evolution with the protein being observed in simple animals such as *T. adhaerens* up to complex mammals such as *H. sapiens*. **C)** Schematic describing the HIF-based O_2_ sensing and signaling system in mammals. Cells sense adequate O_2_ levels (physoxia, shaded blue) via O_2_-dependent proline hydroxylation of HIF-1α leading to subsequent degradation. Under hypoxia (shaded pink), HIF-1α translocates to the nucleus and binds to HIF-1β. This transcription complex binds to DNA and upregulates the transcription of hypoxia response elements (HRE) to restore O_2_ homeostasis.

Iron in PHD2 is coordinated to a 2-oxoglutarate (2OG) cofactor, two histidines and an aspartate from the protein.^[15]^ Under physoxia (3-7.4 % O_2_,^[16]^ region shaded blue in **Fig. 1C**), PHD2 binds co-substrate O_2_ at the iron-center to hydroxylate specific proline residues of HIF-1α.^[17,18]^ Hydroxylation of HIF-1α creates a binding site for the Von Hippel Lindau protein (pVHL) which leads to polyubiquitination and subsequent degradation by the E3 ligase complex.^[19,20]^ Being O_2_-limited, hydroxylation of HIF-1α by PHD2 slows down under hypoxia (<2% O_2_,^[16,21]^ region shaded pink in **Fig. 1C**), leading to its accumulation in the cytosol.^[8]^ Cytosolic HIF-1α translocates to the nucleus and forms a transcription complex with HIF1-β.^[22]^ The transcription complex then binds to hypoxia response gene elements (HRE) to initiate the transcription of 70 genes that modulate physiology via changes in metabolism, angiogenesis, and erythropoiesis to allow for hypoxia adaptation and O_2_ homeostasis restoration.^[6,23,24]^ These hypoxia activated genes show significant upregulation in several disease types such as cancer and Alzheimer’s.^[25–28]^ Consequently, the HIF-signaling pathway has emerged as a prominent therapeutic target for the treatment of these diseases.

Given the therapeutic relevance, approaches that inhibit downstream HIF-signaling pathways have been explored extensively.^[25–30]^ A radically different approach would be to activate PHD2-like O_2_ sensors such that they enhance HIF-1α hydroxylation for subsequent degradation despite hypoxia. A recent computational study suggested diffusion limited O_2_ transport to the active site of PHD2 as the basis for the slow reaction of PHD2 with O_2_.^[31]^ In this study, we use structure-guided computational modeling and protein design approaches to improve O_2_ delivery to PHD2 catalytic core as a pathway to activate HIF-1α hydroxylation. More specifically, we focus on a gas tunnel and critical residues therein that limit the flow of O_2_ to PHD2’s catalytic iron core. We show that rational and systematic modification of these gas tunnel residues can open the constriction topology of PHD2’s gas tunnel and enhance the catalytic efficiency *k*_cat_/*K*_M_(O_2_) from 90 ± 9 M^-1^ s^-1^ for WT PHD2 to 830 ± 40 M^-1^ s^-1^ for the most effectively designed mutant. This mutant involves the substitution of two bulky tryptophan residues in PHD2’s gas tunnel to smaller phenylalanine residues. Furthermore, transfection of plasmids that express these designed PHD2 variants result in a 2-fold reduction of HIF-1α levels in HEK-293T mammalian cells as compared to WT PHD2 under hypoxic conditions (1% O_2_). The impact is further amplified in downstream HIF signaling pathways with 6-fold, 3-fold, and 1.8-fold reduction in the levels of hypoxia response transcripts CA9, HK2, and ADM, respectively. Overall, this study brings forth the use of activated PHD2 as a pathway to manipulate hypoxia signaling in cells and also provides insights into the basis of O_2_ sensing by non-heme iron-based proteins.

## Results and Discussion

### Computation-guided rational design of PHD2’s gas tunnel

We note that the iron atom in PHD2 is buried in the protein’s macromolecular structure such that a tunnel would be required for O_2_ gas to reach the iron core. Such O_2_-delivering tunnels have also been structurally and biochemically characterized for various heme-based O_2_ transporting proteins and gas sensors, such as H-NOX and myoglobin.^[32,33]^ To identify possible O_2_ transport tunnels connecting the catalytic iron-center to the surface of PHD2, we used CAVER^[34]^ – a software tool that visualizes and analyzes tunnels and channels in protein structures. We used the crystal structure of PHD2 complexed with a HIF-1α peptide mimic (PDB: 5L9B) for these analyses since PHD2 is known to bind O_2_ only after its association with HIF-1α.^[35],[36]^ The CAVER program identified a single narrow tunnel created by the interface between PHD2 and the HIF-1α peptide (**Fig. 2A**). The tunnel is mainly comprised of PHD2 residues with bulky non-polar characteristics (I256, W258, M299, I327, and W389) along with some polar residues (Q239, R252, and T387) (**Fig. 2B**). Two residues (P564 and Y565 colored teal, **Fig. 2B**) from the HIF-1α peptide mimic also contributed to forming the tunnel. The predominant non-polar nature of this tunnel aligns with its function as the path for non-polar O_2_ gas. Our analysis of PHD2 crystal structure identified a bottleneck (labeled BN, **Fig. 2B**) in the gas tunnel that was mainly formed by residues M299 and W389 in PHD2 and P564 in HIF-1α. We hypothesized that the presence of large amino acid residues like Met, Trp, Ile, and Arg in PHD2’s gas tunnel could restrict O_2_ transport to PHD2’s active site which may ultimately determine its O_2_ sensing capability and hydroxylation activity. To investigate this possibility, we undertook a computation-and structure-guided rational design approach which involved mutating residues within PHD2’s gas tunnel and investigating its impact on tunnel dynamics and constriction topology.

**Figure 2.**
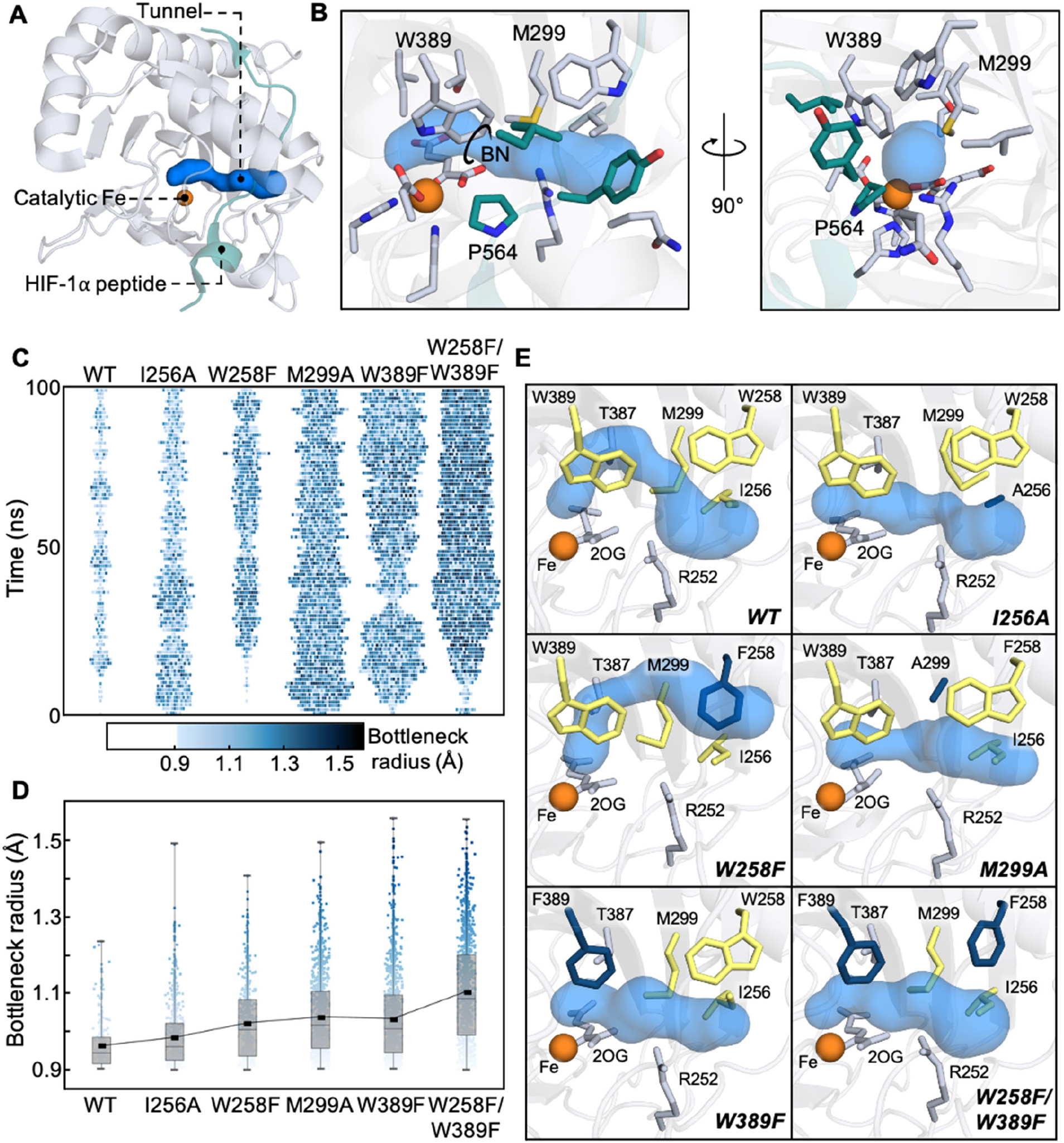
**A)** Full-view and **B)** zoomed-in crystal structure of PHD2 and HIF peptide (PDB: 5L9B) with CAVER computed tunnel. Tunnel is shown in blue with lining residues from PHD2 and HIF peptide shown in gray and teal, respectively. Bottleneck of the tunnel is represented as a circle and labelled BN. **C)** Heat map depicting the frequency and bottleneck radius of the primary tunnel in PHD2 variants from three concatenated 100 ns MD trajectories. The width of lane gives a time-localized estimate of the duration for which the primary tunnel stays open and the color of each block represents the bottleneck radius of each observed tunnel. **D)** Impact of mutations on bottleneck radius (BR) from three concatenated 100 ns MD trajectories. Box plot reveals BR distribution (0, 25, 50, 75 and 100% quantile) for each variant. Trend line connects the average BR across PHD2 variants. **E)** Representative structures from MD simulations of PHD2 variants. The CAVER computed tunnels are shown in blue. WT tunnel forming residues that were targets of our rational design studies are shown in yellow in each panel. Residues that were mutated are shown in dark blue. Residues that potentially constricted the tunnel but were not targets of mutagenesis are shown in gray.

Target residues for our rational design study were selected by assessing multiple criteria such as positioning and orientation of the gas tunnel lining residues, their propensity to obstruct access to the iron core and their conservation in 1844 homologous PHD2 sequences. To begin with, we selected W258 and W389 for our studies as tryptophan has been previously shown to function as gates within tunnels to regulate substrate access and enzyme activity.^[33,37–39]^ W258 is positioned at the entrance to the tunnel, and W389 is positioned directly in front of the iron-center which makes these residues potential entry/exit gates for the O_2_ gas. Additionally, both residues are highly conserved among PHDs (**Fig. S2A,C**) indicating that they may contribute to tuning PHDs’ O_2_ sensing capability across organisms. We also selected M299 as a target residue for mutagenesis due to its role in forming the bottleneck with W389. Methionine is the most conserved amino acid at the 299 position, but other large amino acids such as histidine and glutamine are also observed in other PHDs (**Fig. S2B**). We selected I256 as the fourth residue for rational design studies as it partially obstructs the tunnel entrance along with W258 and is mostly conserved among PHDs (**Fig. S2A**). R252 was not considered as a target residue as it forms a salt bridge with D254 which has been shown to be crucial for HIF-1α binding and hydroxylation activity in PHD2.^[40]^ Lastly, T387 was also excluded as it forms an H-bond with iron’s aqua ligand and mutations to this residue can change the modality of O_2_ binding to iron.^[41]^

Next, we subjected WT PHD2 and mutants of target residues (I256A, W258A, W258F, M299A, W389A, and W389F) to molecular dynamics (MD) simulations to investigate how these mutations would affect protein structure and tunnel dynamics. MD simulations of W258A and W389A PHD2 variants demonstrated significantly altered placement of the HIF-1α peptide mimic and were excluded from further analysis (**Fig. S3-4)**. Instead, we included W258F/W389F PHD2 variant that replaces both bulky tryptophan residues in the tunnel with smaller phenylalanine in subsequent investigations. MD trajectories of WT and selected variants were analyzed using CAVER to understand how individual mutations affected the constriction topology and dynamics of the highest-ranked tunnel, which is also the one identified in the crystal structure. Some other tunnels were observed as well, but they were present minimally leading us to conclude that they did not contribute significantly to O_2_ transport. To visualize how mutations could impact O_2_ transport, we tracked the presence of the highest-ranked O_2_ tunnel within three concatenated 100 ns MD trajectories (3000 total frames) for each PHD2 variant (**Fig. 2C**). Each data point in the plot indicates that the tunnel was found to be present in the MD frame or open for O_2_ transport at that instant, and its color corresponds to its bottleneck radius. The absence of data point corresponds to the tunnel being absent or closed for O_2_ transport. We binned all data points into 1 ns interval along the time axis such that points corresponding to an open tunnel within that 1 ns interval are spread out horizontally. Such a representation eliminates overlapping points that could arise from limited resolution of the plot in the vertical direction. At the same time, the horizontal width of each lane gives a time-localized estimate of the duration for which the primary tunnel stays open as the simulation progresses.

We also computed tunnel statistics for parameters such as the percentage of the total frames showing the primary tunnel, their bottleneck radius (BR), length, and curvature to quantify how the mutations affected the tunnel (**Table 1**). It is easy to see that the first lane (**Fig. 2C**) corresponding to WT PHD2 is characterized by a low density of points and is narrow in width indicating that the tunnel is mostly closed and opens intermittently for short durations to severely limit O_2_ transport. Furthermore, the tunnel when present in WT PHD2 possesses tighter bottlenecks (avg. BR of 0.95 ± 0.05 Å, Fig. 2D). The next two lanes (**Fig. 2C-D**) for I256A and W258F mutants reveal higher point densities and sample wider bottlenecks suggesting potential for improved O_2_ transportation. The last three lanes (**Fig. 2C-D**) corresponding to M299A, W389F and W258F/W389F variants reveal tunnels that stay mostly open (present in 58 ± 19%, 61 ± 15% and 76.2 ± 9.0% of simulation frames, respectively) and sample tunnel topologies with the widest bottlenecks. To gain structural insights into these observations, we compare the highest-ranked tunnel calculated by CAVER using a representative MD frame from each PHD2 variant. Beginning with the MD-simulated WT PHD2, we observe that its gas tunnel (top left panel, **Fig. 2E**) takes a rather curved path to the iron core as compared to the tunnel calculated from the crystal structure (**Fig. 2B**). In the majority of MD frames that contain this gas tunnel, the bottleneck is formed by M299 and 2OG with either R252 or W389. Mutation of I256 to Ala in the I256A variant creates space for bulky M299 to take an alternate conformation (top right panel, **Fig. 2E**) such that the methionine is no longer the bottleneck residue and affords a higher propensity for the tunnel to stay open. Mutating W258 to Phe in the W258F variant modifies the tunnel path at the entry location (middle left panel, **Fig. 2E**) such that O_2_ can take a shorter and less curved path to the iron-center. In the M299A variant, the 299 residue is no longer a bottleneck and O_2_ can take a direct path to the iron-center (middle left panel, **Fig. 2E**). In the W389F and the W258F/W389F variants, the constriction between the 299 and 389 residues are significantly relaxed allowing the tunnel to remain open and offers O_2_ the shortest path to the iron-center (bottom panels, **Fig. 2E**). Overall, our computational protein design studies reveal that it is possible to modify gas tunnel architecture with features that can enhance O_2_ transport to the PHD2 active site.

**Table 1.** CAVER statistics of PHD2 tunnel variants. Reported averages, standard deviations (SD), and average deviations (AD) are across three independent 100 ns MD trajectories.

| Variant | % of frames with primary tunnel | Avg. BR ± AD (Å) | Avg. Length ± AD (Å) | Avg. Curvature ± AD (Å) |
| --- | --- | --- | --- | --- |
| WT | 12 ± 12 | 0.95 ± 0.05 | 19 ± 4 | 1.5 ± 0.3 |
| I256A | 30.1 ± 2.6 | 0.98 ± 0.08 | 18 ± 2 | 1.4 ± 0.2 |
| W258F | 24 ± 10 | 0.97 ± 0.07 | 17 ± 3 | 1.4 ± 0.2 |
| M299A | 58 ± 19 | 1.0 ± 0.1 | 17 ± 2 | 1.3 ± 0.1 |
| W389F | 61 ± 15 | 1.0 ± 0.1 | 15 ± 2 | 1.2 ± 0.2 |
| W258F/W389F | 76.2 ± 9.0 | 1.1 ± 0.1 | 15 ± 2 | 1.2 ± 0.1 |

### Spectroscopic investigation of PHD2 tunnel variants

To experimentally probe the impact of selected mutations on PHD2’s biochemical activity, we expressed and purified WT PHD2 and its I256A, W258F, M299A, W389F and W258F/W389F variants (**Fig. S6**). We characterized the overall protein fold and stability of WT and designed PHD2 variants using circular dichroism (CD) and thermal shift assays (TSA) (**Fig. S7 and S8**, respectively). CD studies showed that the designed variants resembled WT PHD2 in their overall secondary structure. At the same time, TSA data showed that the variants, albeit showing lesser melting temperature than WT, were quite stable at physiological temperature. Next, we used UV-Vis spectroscopy to investigate active-site assembly and binding of HIF-1α peptide mimic to WT PHD2 and a select set of designed variants identified through simulations to have the most open gas tunnels (M299A, W389F, and W258F/W389F PHD2). Non-heme iron hydroxylases exhibit a broad metal-to-ligand-charge-transfer (MLCT) band between Fe^II^ center and the bidentate-coordinated 2OG ligand centered around 475-550 nm. This spectral feature is sensitive to changes in the primary coordination sphere of iron and has been used to study substrate binding and coordination changes in 2OG-dependent non-heme iron enzymes.^[42–45]^ Upon the addition of Fe^II^ and 2OG to WT PHD2, we observed the formation of an MLCT band with a peak centered at around 521 nm (**Fig. 3A**) that can be assigned to a 6c aqua-bound Fe^II^ species, similar to the iron complex previously characterized in taurine dioxygenase (TauD), prolyl-4-hydroxylase (P4H), viral collagen prolyl hydroxylase (vCPH), and clavaminate synthase (CS).^[44,46–48]^ Similar spectral features including λ_max_ and molar absorptivity of the MLCT band were observed for M299A, W389F, and W258F/W389F PHD2 suggesting that the mutations in the gas tunnel of PHD2 did not impact iron binding and the primary coordination sphere (**Fig. S9**). Next, we added the HIF-1α peptide mimic to the 6*c* Fe^II^-2OG complex, which caused the MLCT to blue-shift by 10 nm indicative of loss of the axial aqua ligand and formation of an O_2_ reactive 5*c* Fe^II^-2OG-HIF complex as has been observed in other non-heme iron hydroxylases.^[44,49,50]^ We observed similar spectral transition in M299A, W389F, and W258F/W389F PHD2 suggesting that the tunnel mutations did not affect PHD2’s ability to bind HIF-1α (**Fig. S9**).

**Figure 3.**
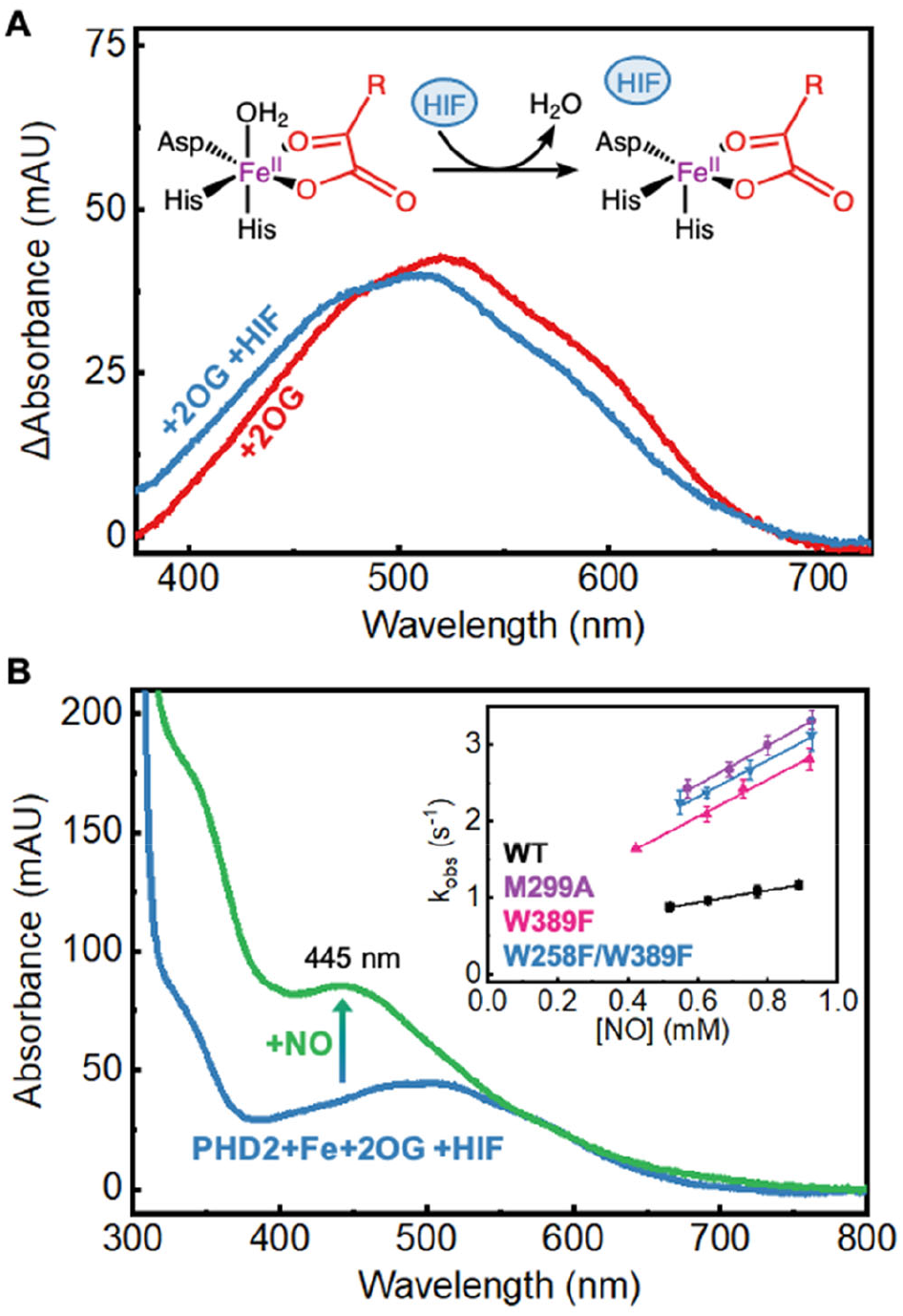
**A)** UV-Vis difference spectra of WT PHD2 active-site assembly and MLCT perturbation. 2OG-bound PHD2 exhibits a broad MLCT centered at 521 nm (red). Binding of the HIF-1α peptide (HIF) forces the loss of an aqua ligand resulting in a 10 nm blue-shift of the MLCT (blue). **B)** UV-Vis spectra of iron nitrosyl formation in WT PHD2 and determination of *k*_on_(NO) for PHD2 variants (inset). Upon exposure to NO, PHD2 forms an iron nitrosyl complex which exhibits a distinct UV-Vis signature at 445 nm (green). Monitoring the kinetics of nitrosyl formation (*k*_obs_) at varying NO concentrations leads to the determination of *k*_on_(NO) for PHD2 variants (inset).

To experimentally confirm enhanced gas diffusion through engineered tunnels to the iron center of these mutants, we employed UV-Vis stopped flow kinetics and measured the association rates (*k*_on_) of these mutants with nitric oxide (NO) that was used as a surrogate for O_2_. While O_2_ is the natural co-substrate, NO forms a stable iron nitrosyl species characterized by an intense peak at 445 nm (**Fig. 3B, Fig. S10**) that allows stopped-flow UV-Vis spectroscopy based kinetic measurements of diatomic gas binding to the iron center.^[51,52]^ Using this, we determined *k*_on_(NO) as the slope of observed NO-binding rates (*k*_obs_) of WT PHD2 and its variants plotted against total NO concentrations used (inset, **Fig. 3B**). In turn, we find that WT PHD2 exhibits a *k*_on_(NO) of 0.771 ± 0.018 mM^-1^ s^-1^, while the M299A, W389F and W258F/W389F variants exhibit a ∼3.5x increase in *k*_on_(NO) of 2.55 ± 0.13 mM^-1^ s^-1^, 2.41 ± 0.08 mM^-1^ s^-1^ and 2.39 ± 0.15 mM^-1^ s^-1^, respectively, confirming enhanced diatomic gas diffusion in the tunnel engineered mutants.

### Catalytic hydroxylation activity of PHD2 tunnel variants

Next, we performed an activity screen of WT PHD2 and all of the computationally designed variants by monitoring the rate of conversion of a HIF-1α peptide mimic to the hydroxylated peptide using matrix assisted laser desorption/ionization time-of-flight (MALDI-TOF) mass spectrometry (**Fig. S11**).^[53]^ Under atmospheric O_2_ levels (256 μM or 21% O_2_), WT PHD2 exhibits a hydroxylation rate of 0.016 ± 0.001 s^-1^ (**Fig. 4A**). While I256A PHD2 displayed a tunnel with slightly improved access in the simulation, the mutant exhibits hydroxylation activity very similar to WT. The W258F variant shows a 1.25-fold increase in hydroxylation rate suggesting W258’s role as a tryptophan gate obstructing the O_2_ gas tunnel at its entrance. The M299A PHD2 mutant exhibited a 3-fold reduced hydroxylation rate indicating that the M299 residue plays a functional role in hydroxylation catalysis beyond simply controlling/limiting O_2_ access. We note that M299 residue participates in hydrophobic sulfur-ring bonds with W389 and W258 residues forming an aromatic-methionine bridge motif (**Fig. S12**) that are ubiquitous in proteins offering structural stability and defense against oxidation of the aromatic groups by reactive species.^[54–56]^ While the M299A mutation disrupts this functionally important motif, they are retained in W258F and W389F PHD2 mutants (**Fig. S12**). Mutation of the W389 residue to phenylalanine in the W389F variant reveals a 2-fold enhancement in hydroxylation catalysis rate further affirming that the bottleneck forming W389 residue could be the major tryptophan gate in PHD2’s gas tunnel. Ultimately, tandem mutations to both tryptophan gates in the W258F/W389F variant lead to the highest 2.5-fold enhancement in hydroxylation rate and is the most promising mutant from the screen.

**Figure 4.**
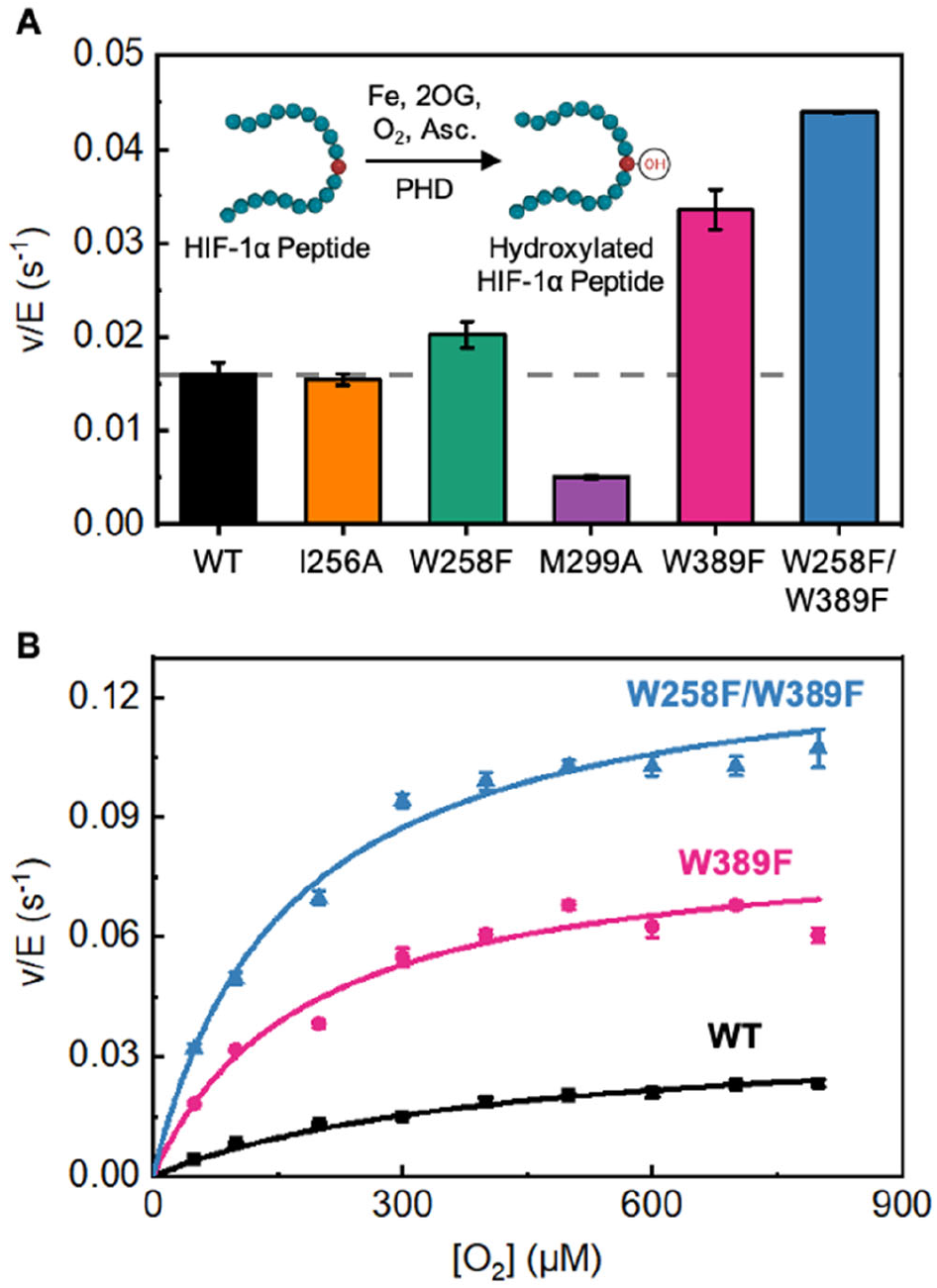
**A)** Hydroxylation activity screen for PHD variants at ambient O_2_ (21% or 256 µM O_2_). Hydroxylation activity was determined by monitoring the conversion of a HIF-1α peptide mimic via MALDI-TOF MS. Averages and S.D. are from three independent experiments. **B)** Steady-state kinetics as a function of varied O_2_ concentrations for PHD variants. Averages and S.D. for each data point are from two independent experiments.

To investigate whether these tryptophan gates exert control on PHD2’s hydroxylation catalysis through an O_2_ dependent pathway, we conducted steady-state kinetic studies at varying O_2_ concentrations using WT PHD2 and the highest activity variants, W389F and W258F/W389F. Fitting the data to the Michaelis-Menten model, we calculate a *K*_M_(O_2_) of 415 ± 36 µM, a catalytic first-order rate constant (*k*_cat_) of 0.037 ± 0.002 s^-1^, and catalytic efficiency (*k*_cat_/*K*_M_(O_2_)) of 90 ± 9 M^-1^ s^-1^ for WT PHD2 (black curve, **Fig. 4B**) which matches previously reported^[11]^ values (**Table 2, Fig. S13**). The W389F PHD2 variant reveals a 2.5-fold reduced *K*_M_(O_2_) (183 ± 16 µM) and a 2-fold enhanced *k*_cat_ (0.085 ± 0.005 s^-1^) yielding a 5-fold enhancement in catalytic efficiency (470 ± 19 M^-1^ s^-1^) relative to WT PHD2. While the *K*_M_(O_2_) of the W258F/W389F PHD2 variant (161 ± 17 µM) is comparable to the W389F PHD2 variant, it has a 3.6-fold enhanced *k*_cat_ (0.134 ± 0.005 s^-1^) compared to WT PHD2. Overall, W258F/W389F PHD2 exhibited a 9-fold higher *k*_cat_/*K*_M_(O_2_) as compared to WT PHD2 suggesting that engineering of PHD2’s gas tunnel can indeed enhance its HIF-1α hydroxylation performance. The increase in *k*_cat_ values of W389F and W258F/W389F PHD2 variants suggests that catalytic steps past O_2_ association are enhanced as well. We note that oxidative decarboxylation of 2OG at the active site produces CO_2_ gas. Akin to O_2_ entry, CO_2_ egress from the catalytic pocket has to occur through a gas tunnel and can impact product release related contributions to *k*_cat_. In turn, we simulated activated PHD2 variants with the iron-center in the form of the ferryl-intermediate to evaluate the tunnel characteristics for CO_2_ egress. The simulations revealed that the engineered mutants offer a tunnel that is more accessible for CO_2_ release from the active site relative to the WT and can contribute to enhanced *k*_cat_ (**Fig. S14** and **Table S2**).

**Table 2.** O_2_ dependent kinetic constants of WT and tunnel engineered PHD2 variants.

| Variant | $K_M(O_2)$<br>($\mu\text{M}$) | $k_{cat}$<br>($\text{s}^{-1}$) | $k_{cat}/K_M(O_2)$<br>($\text{M}^{-1} \text{s}^{-1}$) |
| --- | --- | --- | --- |
| WT | $415 \pm 36$ | $0.037 \pm 0.002$ | $90 \pm 9$ |
| W389F | $183 \pm 16$ | $0.085 \pm 0.005$ | $470 \pm 19$ |
| W258F/W389F | $161 \pm 17$ | $0.134 \pm 0.005$ | $830 \pm 40$ |

### Impact of PHD2 tunnel variants on hypoxia signaling in cells

The results from biochemical assays encouraged us to investigate the impact of designed PHD2 variants on cellular HIF-1α levels under hypoxic conditions. PHD2 senses the presence of O_2_ in cells by catalyzing hydroxylation of HIF-1α, which primes the latter for proteasomal degradation. The relatively low O_2_ levels under hypoxia inhibits PHD2 resulting in cellular accumulation of HIF-1α. To investigate if designed PHD2 variants modulate cellular HIF-1α levels, we transfected HEK-293T cells with plasmids expressing WT, W389F, and W258F/W389F PHD2 variants and incubated them under atmospheric O_2_ levels (21% O_2_) and hypoxic (1% O_2_) conditions (**Fig. 5A**). We used western blot to monitor intracellular HIF-1α and PHD2 levels with α-tubulin as the loading control. At 21% O_2_, transfection of both WT and PHD2 variants led to undetectable HIF1α levels (**Fig. S15**).

**Figure 5.**
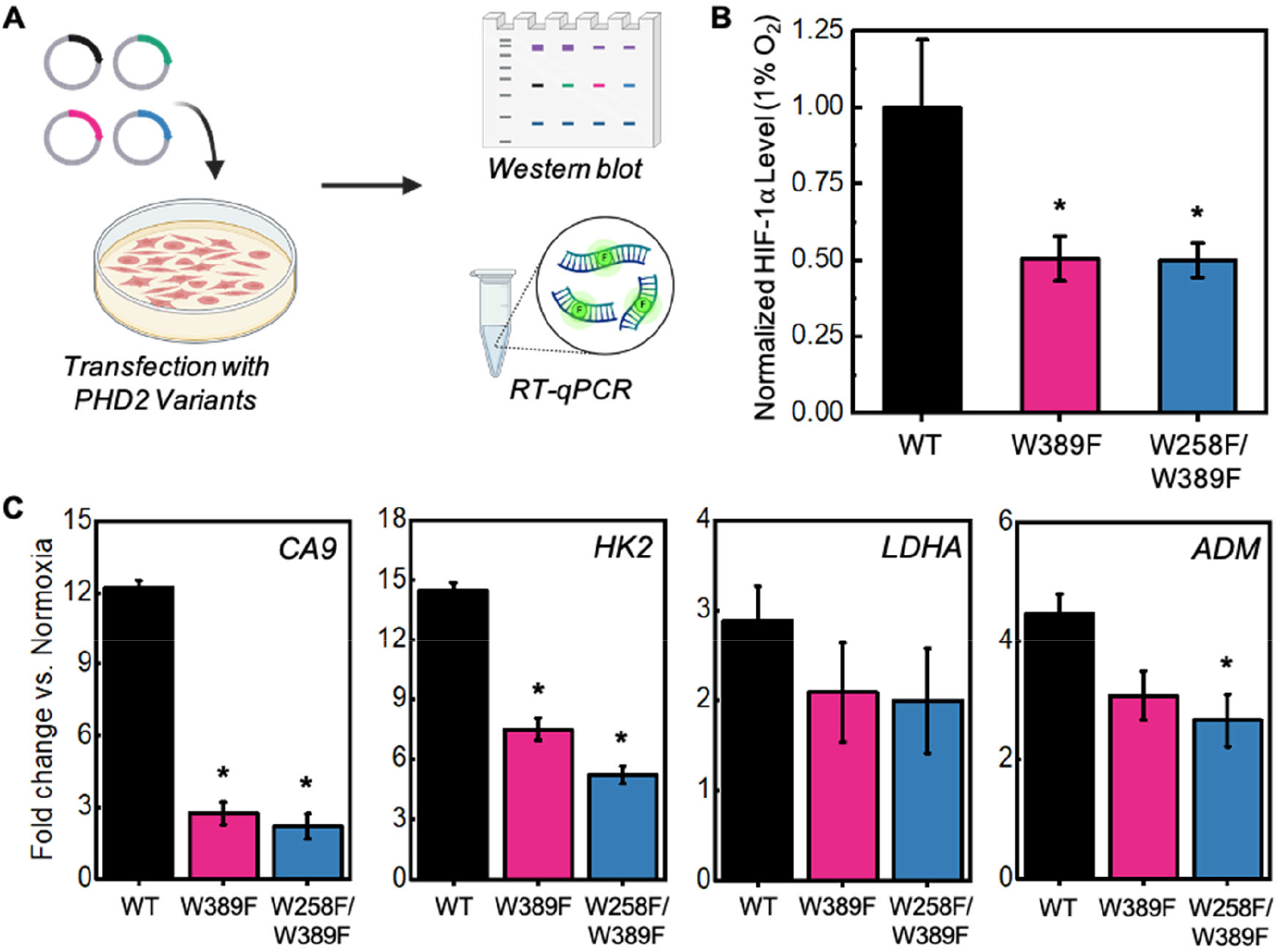
**A)** Schematic of hypoxia signaling experiments. HEK-293T cells were transfected with WT and PHD2 variants and incubated under hypoxia (1% O_2_). HIF-1α levels were determined using western blot. Downstream transcripts were quantified using RT-qPCR. **B)** HIF-1α protein levels from western blot. Values were quantified in ImageJ and normalized by WT PHD2 using the equation: *normalized HIF-1α = HIF-1α(PHD2 variant/WT PHD2)*. Averages and S.D. are from three independent experiments. Asterisks denote statistical significance compared to WT PHD2 (*p < 0.05). **C)** RT-qPCR results for the quantification of CA9, HK2, LDHA, and ADM transcripts after transfection with PHD2 variants. Averages and S.D. are from three independent experiments. Asterisks denote statistical significance compared to WT PHD2 (*p < 0.05).

Incubation under hypoxia (1% O_2_) allowed for HIF-1α accumulation in cells, so any differences in HIF-1α levels between WT PHD2 and engineered variants could be quantified (**Fig. S15**). HIF-1α levels were normalized against a ratio of PHD2 variant to WT PHD2 to ensure that any variation in the levels of transfection and protein expression did not skew quantification. These studies revealed that the transfection of plasmids expressing W389F and W258F/W389F PHD2 variants demonstrated a 2-fold reduction in HIF-1α protein levels as compared to WT PHD2, suggesting that the expression of PHD2 variants enhanced hydroxylation and proteasomal degradation of endogenous HIF-1α under hypoxia conditions (**Fig. 5B**).

Due to HIF-1α’s role as a transcription factor, we were also interested in how PHD2 variants impact downstream transcriptional responses. Using RT-qPCR, we investigated relative changes of several common hypoxia response genes involved in pH balance, glycolysis and angiogenesis. First, we probed for differences in carbonic anhydrase IX (CA9) mRNA levels since CA9 plays an important role in maintaining cellular pH balance and is a promising marker and target for anticancer therapy.^[57,58]^ Cells transfected with engineered PHD2 variant plasmids revealed a significant 6-fold reduction in mRNA levels corresponding to the carbonic anhydrase IX (CA9) gene as compared to those transfected with WT PHD2 (**Fig. 5C**).

Next, we probed for differences in mRNA levels of glycolytic enzymes hexokinase 2 (HK2) and lactate dehydrogenase A (LDHA) that are upregulated under hypoxia to enhance anaerobic energy production. Our results showed that the average levels of both LDHA and HK2 transcripts were reduced upon the expression of PHD2 variants compared to WT PHD2. While the change in LDHA transcript levels did not reach statistical significance due to measurement variations, HK2 transcript levels demonstrated 2-3 fold reduction in cells transfected with designed PHD2 variants compared to WT PHD2 (**Fig. 5C**). Finally, we investigated adrenomedullin (ADM) transcript levels that trigger vasodilation and angiogenesis phenotypes under hypoxia. The average levels of ADM transcript were reduced in cells expressing PHD2 variants compared to WT PHD and the cells transfected with the W258F/W389F variant plasmid yielded a significant 1.8-fold reduction compared to WT PHD2 (**Fig. 5C**). Overall, these studies demonstrate that activated PHD2 variants offer a new pathway to reprogram hypoxia responses and HIF signaling in cells.

## Conclusion

Biocatalysts play important roles in physiology, however, their engineered forms have been rarely used to enhance cellular function. Several challenges including the efficient delivery of the biocatalyst along with all the required co-substrates to the cell and the need to ensure that the engineered biocatalyst retains all protein-protein interactions (PPI) pertaining to their function have limited such possibilities. In this work, we have circumvented these challenges and have demonstrated the ability to enhance HIF-1α hydroxylation and degradation in human kidney cells using engineered PHD2 mutants. We note that hypoxia-induced HIF-1α and its downstream transcripts accumulate in various disease types, including cancer and Alzheimer’s and numerous small-molecule and gene-editing methodologies that target HIF-1α have been developed, some of which are currently in clinical trials.^[29,30]^ While downstream inhibition of HIF signaling pathways are being aggressively pursued, we have achieved the same by activating PHD2, a biocatalyst that is upstream to the transcription factor. In all, our work represents a new modality in which we activate enzymatic sensors upstream to the transcription factor as compared to conventional strategies of inhibiting downstream signaling cascades. Beyond presenting a novel application of engineered biocatalysts, our studies also represent a systematic investigation into how gas tunnels in PHD2-like enzymatic sensors can be engineered to control their O_2_ reactivity and catalytic efficiency. Overall, this work elucidates how non-heme iron proteins utilize gas tunnels to tune their O_2_ sensing capability to match physiological needs as well as presents a novel application of engineered biocatalysts in the realm of cellular biology and hypoxia signaling.

## Supporting information

Supp Info_gas tunnel hypoxia signaling

## Supporting Information

The authors have cited additional references within the Supporting Information.^[59–70]^

## Acknowledgements

Bhagi-Damodaran Lab acknowledges funds from Regents of the University of Minnesota, American Cancer Society and RCSA Cottrell Foundation for supporting this research. The development of O_2_-dependent assays in this work was funded by NIH NIGMS grant #R35GM138277. Chen lab acknowledge their funding resources NIH Grant (R35GM124896) and NSF Grant CHE-1753154.

## Notes

### Competing Interest Statement

The authors have declared no competing interest.

### Summary of Updates

More datasets have been added including spectroscopic and kinetics studies as well as downstream signaling data.

## References

[1] F. W. Outten, E. C. Theil, Antioxid. Redox Signal. 2009, 11, 1029–1046.

[2] P. Brzezinski, A. Moe, P. Ädelroth, Chem. Rev. 2021, 121, 9644–9673.

[3] M. R. Dent, B. R. Weaver, M. G. Roberts, J. N. Burstyn, J. Bacteriol. 2023, 205, e00332–22.

[4] F. Zhong, G. P. Lisi, D. P. Collins, J. H. Dawson, E. V. Pletneva, Proc. Natl. Acad. Sci. 2014, 111, DOI 10.1073/pnas.1317173111.

[5] N. J. Hoque, E. E. Weinert, Curr. Opin. Microbiol. 2023, 76, 102396.

[6] C. J. Schofield, P. J. Ratcliffe, Nat. Rev. Mol. Cell Biol. 2004, 5, 343–354.

[7] R. K. Bruick, S. L. McKnight, Science 2001, 294, 1337–1340.

[8] A. C. R. Epstein, J. M. Gleadle, L. A. McNeill, K. S. Hewitson, J. O’Rourke, D. R. Mole, M. Mukherji, E. Metzen, M. I. Wilson, A. Dhanda, Y.-M. Tian, N. Masson, D. L. Hamilton, P. Jaakkola, R. Barstead, J. Hodgkin, P. H. Maxwell, C. W. Pugh, C. J. Schofield, P. J. Ratcliffe, Cell 2001, 107, 43–54.

[9] L. Brewitz, A. Tumber, C. J. Schofield, J. Biol. Chem. 2020, 295, 7826–7838.

[10] H. Tarhonskaya, A. P. Hardy, E. A. Howe, N. D. Loik, H. B. Kramer, J. S. O. McCullagh, C. J. Schofield, E. Flashman, J. Biol. Chem. 2015, 290, 19726–19742.

[11] T. Liu, M. I. Abboud, R. Chowdhury, A. Tumber, A. P. Hardy, K. Lippl, C. T. Lohans, E. Pires, J. Wickens, M. A. McDonough, C. M. West, C. J. Schofield, J. Biol. Chem. 2020, 295, 16545–16561.

[12] M. T. Williams, E. Yee, G. W. Larson, E. A. Apiche, A. Rama Damodaran, A. Bhagi-Damodaran, Curr. Opin. Chem. Biol. 2023, 76, 102331.

[13] J. S. Scotti, I. K. H. Leung, W. Ge, M. A. Bentley, J. Paps, H. B. Kramer, J. Lee, W. Aik, H. Choi, S. M. Paulsen, L. A. H. Bowman, N. D. Loik, S. Horita, C. Ho, N. J. Kershaw, C. M. Tang, T. D. W. Claridge, G. M. Preston, M. A. McDonough, C. J. Schofield, Proc. Natl. Acad. Sci. 2014, 111, 13331–13336.

[14] K. Lippl, A. Boleininger, M. McDonough, M. I. Abboud, H. Tarhonskaya, R. Chowdhury, C. Loenarz, C. J. Schofield, Hypoxia 2018, Volume 6, 57–71.

[15] M. A. McDonough, V. Li, E. Flashman, R. Chowdhury, C. Mohr, Proc. Natl. Acad. Sci. 2006, 103, 9814–9819.

[16] S. R. McKeown, Br. J. Radiol. 2014, 87, 20130676.

[17] W. He, S. Batty-Stuart, J. E. Lee, M. Ohh, J. Mol. Biol. 2021, 433, 167244.

[18] E. Flashman, E. A. L. Bagg, R. Chowdhury, J. Mecinović, C. Loenarz, M. A. McDonough, K. S. Hewitson, C. J. Schofield, J. Biol. Chem. 2008, 283, 3808–3815.

[19] J.-H. Min, H. Yang, M. Ivan, F. Gertler, W. G. Kaelin, N. P. Pavletich, Science 2002, 296, 1886–1889.

[20] J. Lisztwan, G. Imbert, C. Wirbelauer, M. Gstaiger, W. Krek, Genes Dev. 1999, 13, 1822–1833.

[21] D. Xie, T. L. King, A. Banerjee, V. Kohli, E. L. Que, J. Am. Chem. Soc. 2016, 138, 2937–2940.

[22] S. M. Wood, J. M. Gleadle, C. W. Pugh, O. Hankinson, P. J. Ratcliffe, J. Biol. Chem. 1996, 271, 15117–15123.

[23] B.-H. Jiang, E. Rue, G. L. Wang, R. Roe, G. L. Semenza, J. Biol. Chem. 1996, 271, 17771–17778.

[24] R. H. Wenger, D. P. Stiehl, G. Camenisch, Sci. STKE 2005, 2005, DOI 10.1126/stke.3062005re12.

[25] L. Semenza, J. Clin. Invest. 2013, 123, 3664–3671.

[26] B. Muz, P. de la Puente, F. Azab, A. K. Azab, Hypoxia 2015, 83.

[27] X. Sun, G. He, H. Qing, W. Zhou, F. Dobie, F. Cai, M. Staufenbiel, L. E. Huang, W. Song, Proc. Natl. Acad. Sci. 2006, 103, 18727–18732.

[28] R. March-Diaz, N. Lara-Ureña, C. Romero-Molina, A. Heras-Garvin, C. Ortega-de San Luis, M. I. Alvarez-Vergara, M. A. Sanchez-Garcia, E. Sanchez-Mejias, J. C. Davila, A. E. Rosales-Nieves, C. Forja, V. Navarro, A. Gomez-Arboledas, M. V. Sanchez-Mico, A. Viehweger, A. Gerpe, E. J. Hodson, M. Vizuete, T. Bishop, A. Serrano-Pozo, J. Lopez-Barneo, E. Berra, A. Gutierrez, J. Vitorica, A. Pascual, Nat. Aging 2021, 1, 385–399.

[29] Z. Li, Q. You, X. Zhang, J. Med. Chem. 2019, 62, 5725–5749.

[30] M. Li, H. Xie, Y. Liu, C. Xia, X. Cun, Y. Long, X. Chen, M. Deng, R. Guo, Z. Zhang, Q. He, J. Controlled Release 2019, 304, 204–215.

[31] C. Domene, C. Jorgensen, C. J. Schofield, J. Am. Chem. Soc. 2020, 142, 2253–2263.

[32] R. F. Tilton, I. D. Kuntz, G. A. Petsko, Biochemistry 1984, 23, 2849–2857.

[33] M. B. Winter, M. A. Herzik, J. Kuriyan, M. A. Marletta, Proc. Natl. Acad. Sci. 2011, 108, DOI 10.1073/pnas.1114038108.

[34] E. Chovancova, A. Pavelka, P. Benes, O. Strnad, J. Brezovsky, B. Kozlikova, A. Gora, V. Sustr, M. Klvana, P. Medek, L. Biedermannova, J. Sochor, J. Damborsky, PLoS Comput. Biol. 2012, 8, e1002708.

[35] R. Chowdhury, I. K. H. Leung, Y.M. Tian, M. I. Abboud, W. Ge, C. Domene, F.X. Cantrelle, I. Landrieu, A. P. Hardy, C. W. Pugh, P. J. Ratcliffe, T. D. W. Claridge, C. J. Schofield, Nat. Commun. 2016, 7, 12673.

[36] H. Tarhonskaya, R. Chowdhury, I. K. H. Leung, N. D. Loik, J. S. O. McCullagh, T. D. W. Claridge, C. J. Schofield, E. Flashman, Biochem. J. 2014, 463, 363–372.

[37] L. Biedermannová, Z. Prokop, A. Gora, E. Chovancová, M. Kovács, J. Damborský, R. C. Wade, J. Biol. Chem. 2012, 287, 29062–29074.

[38] S. M. Kim, J. Lee, S. H. Kang, Y. Heo, H.J. Yoon, J.-S. Hahn, H. H. Lee, Y. H. Kim, Nat. Catal. 2022, 5, 807–817.

[39] J. C. Jones, R. Banerjee, K. Shi, H. Aihara, J. D. Lipscomb, Biochemistry 2020, 59, 2946–2961.

[40] R. Chowdhury, M. A. McDonough, J. Mecinović, C. Loenarz, E. Flashman, K. S. Hewitson, C. Domene, C. J. Schofield, Structure 2009, 17, 981–989.

[41] S. Pektas, C. Y. Taabazuing, M. J. Knapp, Biochemistry 2015, 54, 2851–2857.

[42] E. R. Smithwick, R. H. Wilson, S. Chatterjee, Y. Pu, J. J. Dalluge, A. R. Damodaran, A. Bhagi-Damodaran, ACS Catal. 2023, 13, 13743–13755.

[43] K. M. Light, J. A. Hangasky, M. J. Knapp, E. I. Solomon, J. Am. Chem. Soc. 2013, 135, 9665–9674.

[44] M. J. Ryle, R. Padmakumar, R. P. Hausinger, Biochemistry 1999, 38, 15278–15286.

[45] M. L. Neidig, C. D. Brown, K. M. Light, D. G. Fujimori, E. M. Nolan, J. C. Price, E. W. Barr, J. M. Bollinger, C. Krebs, C. T. Walsh, E. I. Solomon, J. Am. Chem. Soc. 2007, 129, 14224–14231.

[46] L. M. Hoffart, E. W. Barr, R. B. Guyer, J. M. Bollinger, C. Krebs, Proc. Natl. Acad. Sci. 2006, 103, 14738–14743.

[47] J. E. Longbotham, C. Levy, L. O. Johannissen, H. Tarhonskaya, S. Jiang, C. Loenarz, E. Flashman, S. Hay, C. J. Schofield, N. S. Scrutton, Biochemistry 2015, 54, 6093–6105.

[48] E. G. Pavel, J. Zhou, R. W. Busby, M. Gunsior, C. A. Townsend, E. I. Solomon, J. Am. Chem. Soc. 1998, 120, 743–753.

[49] E. L. Hegg, A. K. Whiting, R. E. Saari, J. McCracken, R. P. Hausinger, L. Que, Biochemistry 1999, 38, 16714–16726.

[50] J. Zhou, M. Gunsior, B. O. Bachmann, C. A. Townsend, E. I. Solomon, J. Am. Chem. Soc. 1998, 120, 13539–13540.

[51] S. Tian, R. Fan, T. Albert, R. L. Khade, H. Dai, K. A. Harnden, P. Hosseinzadeh, J. Liu, M. J. Nilges, Y. Zhang, P. Moënne-Loccoz, Y. Guo, Y. Lu, Chem. Sci. 2021, 12, 6569–6579.

[52] B. S. Rivard, M. S. Rogers, D. J. Marell, M. B. Neibergall, S. Chakrabarty, C. J. Cramer, J. D. Lipscomb, Biochemistry 2015, 54, 4652–4664.

[53] M. A. Mingroni, V. Chaplin Momaney, A. N. Barlow, I. Jaen Maisonet, M. J. Knapp, in Methods Enzymol., Elsevier, 2023, pp. 363–380.

[54] G. Kim, S. J. Weiss, R. L. Levine, Biochim. Biophys. Acta BBA - Gen. Subj. 2014, 1840, 901–905.

[55] D. S. Weber, J. J. Warren, J. Inorg. Biochem. 2018, 186, 34–41.

[56] D. S. Weber, J. J. Warren, Arch. Biochem. Biophys. 2019, 672, 108053.

[57] S. Pastorekova, R. J. Gillies, Cancer Metastasis Rev. 2019, 38, 65–77.

[58] H.J. Shin, S. B. Rho, D. C. Jung, I.O. Han, E.S. Oh, J.Y. Kim, J. Cell Sci. 2011, 124, 1077–1087.

[59] L. Fu, B. Niu, Z. Zhu, S. Wu, W. Li, Bioinformatics 2012, 28, 3150–3152.

[60] A. Larsson, Bioinformatics 2014, 30, 3276–3278.

[61] G. E. Crooks, G. Hon, J.M. Chandonia, S. E. Brenner, Genome Res. 2004, 14, 1188–1190.

[62] K. Tamura, G. Stecher, S. Kumar, Mol. Biol. Evol. 2021, 38, 3022–3027.

[63] A. Jurcik, D. Bednar, J. Byska, S. M. Marques, K. Furmanova, L. Daniel, P. Kokkonen, J. Brezovsky, O. Strnad, J. Stourac, A. Pavelka, M. Manak, J. Damborsky, B. Kozlikova, Bioinformatics 2018, 34, 3586–3588.

[64] E. F. Pettersen, T. D. Goddard, C. C. Huang, G. S. Couch, D. M. Greenblatt, E. C. Meng, T. E. Ferrin, J. Comput. Chem. 2004, 25, 1605–1612.

[65] P. Li, K. M. Merz, J. Chem. Inf. Model. 2016, 56, 599–604.

[66] W. L. Jorgensen, J. D. Madura, J. Am. Chem. Soc. 1983, 105, 1407–1413.

[67] R. Salomon-Ferrer, A. W. Götz, D. Poole, S. Le Grand, R. C. Walker, J. Chem. Theory Comput. 2013, 9, 3878–3888.

[68] R. Prins, P. Windsor, B. R. Miller, S. Maiden, Dev. Dyn. 2022, 251, 1741–1753.

[69] P. K. Windsor, S. P. Plassmeyer, D. S. Mattock, J. C. Bradfield, E. Y. Choi, B. R. Miller, B. H. Han, Int. J. Mol. Sci. 2021, 22, 2888.

[70] D. R. Roe, T. E. Cheatham, J. Chem. Theory Comput. 2013, 9, 3084–30

