## Supplementary material for "Gas tunnel engineering of prolyl hydroxylase reprograms hypoxia signaling in cells": Supp Info_gas tunnel hypoxia signaling

### Materials and Methods

***Materials:*** All chemicals were purchased from commercial vendors. All primers were purchased from Integrated DNA Technologies (Coralville, IA, USA). The sequence of the HIF-1α peptide mimic was DLDLEMLAPYIPMDDDFQL which is similar to the peptide substrates used in previous studies.[10,18,36] The peptide (99% purity) was synthesized and purchased from Peptide Synthesis Services (University of Minnesota, Internal Service Organization).

***Bioinformatic analysis*****:** To observe sequence conservation at the 256, 258, 299, and 389 residue positions of PHD2, the PHD2 amino acid sequence ($PHD2_{1-426}$) was BLAST searched in the non-redundant protein database. An E-value cutoff of $e^{-9}$ produced 4,966 protein sequences. The sequences were filtered using CD-HIT to remove redundant sequences of 99% similarity.[59] The sequences were further filtered to remove sequences that contained more than 550 amino acids or less than 300 amino acids to ensure the sequences had similar sequence length to PHD2 (426 amino acids). This produced 1844 sequences for the final alignment. The sequences were aligned using the MAFFT alignment algorithm. Alignment results were viewed in AliView, and logos plots were created in WebLogo. [60,61] Phylogenetic trees were constructed in the MEGA 11 software using the neighbor joining method (1000 replicates). [62] Sequences were clustered in groups of 10 to ensure adequate visualization of tree anatomy.

***Tunnel analysis of PHD2 crystal structure*****:** CAVER Analyst 2[63] was used to identify possible $O_2$ pathways in the WT PHD2 crystal structure (PDB: 5L9B). [35] The catalytic iron atom was used as the starting point with a maximum starting distance of 3 Å and a desired radius of 5 Å. The minimum probe radius was set to 0.9 with a shell depth of 4 and a shell radius of 3. The tunnel clustering threshold was set to 3.5.

***Molecular dynamics of PHD2 variants*****:** Molecular dynamics (MD) simulations were used to determine how amino acid mutations affected the proposed tunnels of PHD2 and its variants The starting structure of WT PHD2 bound to 2OG and a HIF-1α peptide mimic was taken from the Protein Data Bank (PDB: 5L9B) and mutations were induced using UCSF Chimera. [35,64] Parameters for the PHD iron-center were determined using the *MCPB.py* module of Amber16. [65] Explicit hydrogen atoms were added to the protein and its mutant structures using the *tleap* module of Amber16. Proteins were solvated in a truncated octahedron unit cell with TIP3P water molecules (neutralized with $Na^+$ ions) with a 12.0 Å solvent buffer between the protein and the closest edge of the unit cell. [66]

The system was simulated with the Amber ff14SB forcefield. The GPU-accelerated *pmemd* code of Amber16 was used to perform all steps of MD.[67] The system was minimized using a seven-step process consisting of 1000 steps of steepest descent minimization followed by 5000 steps of conjugate gradient minimization for each step. For the first step of minimization, all solute heavy atoms were subjected to restraints starting at 10.0 kcal/mol/Å$^2$. Restraints were lowered systematically over the course of each step with the last step having a restraint of 0.0 kcal/mol/Å$^2$. Next, the system was heated linearly from 10.0 K to 298.0 K over 2.0 ns. During heating, all solute atoms were subjected to a restraint of 10.0 kcal/mol/Å$^2$. A maximum temperature of 298.0 K was chosen in an effort to mimic the conditions at which the experimental trials were completed. Equilibration of the system was carried out over 3.5 ns at a constant temperature of 298.0 K. All solute heavy atoms were subjected to restraints of 10.0 kcal/mol/Å$^2$ for the first 0.5 ns, and restraints were decreased geometrically every 0.5 ns until reaching a restraint weight of 0.0 kcal/mol/Å$^2$ for the final 0.5 ns. After equilibration of the system, unrestrained MD was performed at a constant pressure of 1 atm and constant temperature of 298.0 K. The coordinates were saved every 100 ps, and all systems were simulated without restraints for 100 ns in triplicate. Minimization, heating, equilibration, and unrestrained MD is in accordance with previously published works.[68,69] MD analysis (RMSD, b-factor, and distance) was performed using the *cpptraj* module of Amber16.[70] The resulting values for each simulation were averaged for each PHD2 variant and reported. The visualization of the completed trajectories was carried out using PyMOL and UCSF Chimera.[64]

***Tunnel analysis of molecular dynamics trajectories*****:** CAVER Analyst 2[63] was used to identify possible $O_2$ pathways in the in the MD trajectories of PHD2 variants. Each of the triplicate 100 ns trajectories (1000 frames) for each PHD variant (6 variants) were analyzed in this way (18,000 total frames). The catalytic iron atom was used as the starting point with a maximum starting distance of 3 Å and a desired radius of 5 Å. The minimum probe radius was set to 0.9 with a shell depth of 4 and a shell radius of 3. The tunnel clustering threshold was set to 3.5. The tunnel parameters (occupancy, average bottleneck radius, maximum bottleneck radius, length, and curvature) for each simulation were averaged for each PHD2 variant and reported.

***Site-directed mutagenesis*****:** Site-directed mutagenesis was used to create point mutations in PHD2. For recombinant protein expression, $PHD2_{181\text{-}426}$ (wild-type and mutants) was incorporated into pET-28a(+) expression vector with an N-terminal $His_6$ tag.

Site-directed mutagenesis was performed on the wild-type PHD plasmid to create I256A, W258F, M299A, W389F, and W258F/W389F mutations using Phusion site-directed mutagenesis kit (Thermofisher Scientific). The primers (5' → 3') for site-directed mutagenesis were: I256A (forward – GAGGCGATAAGGCCACCTGGATCGA, reverse – GGATGTCCTTGGACGAGTCACTCTTC), W258F (forward – GATAAGATCACCTTCATCGAGGGCAAG, reverse – GCCTCGGATGTCCTTGGACGAGT), M299A (forward – CGGACGAAAGCCGCGGTTGCTTGTTAT, reverse – GCCATTGATTT TGTAGCTGCCCAGCT), W389F (forward – CGCAATAACTGTTTTCTATTTTGATGCAGA, reverse – TACCTTGTAGCATATGCTGGTTGTACTTC). PCR was executed using a standard two-step protocol. Presence of mutations were confirmed using classic Sanger sequencing at University of Minnesota Genomics Center (UMGC). T7 promoter (TAATACGACTCACTATAGGG) and T7 terminator (GCTAGTTATTGCTCAGCGG) were used for sequencing. For mammalian cell transfection, $PHD2_{1\text{-}426}$ (wild-type and mutants) was incorporated into a prk5-HA expression vector. Site-directed mutagenesis was performed on the wild-type PHD2 plasmid to create W389F, and W258F/W389F mutations using Phusion site-directed mutagenesis kit (Thermofisher Scientific). The primers for site-directed mutagenesis were the same as previously mentioned. PCR was executed using a standard two-step protocol. Presence of mutations were confirmed using Nanopore sequencing at Plasmidsaurus.

***Protein expression***: BL-21(DE3) *E. coli* (Thermofisher Scientific) were transformed with pET-28a(+) expression vector containing His6-$PHD_{181\text{-}426}$ (wild-type and mutants). Cells were grown in 2XYT media supplemented with kanamycin (50 µg/mL) at 37 ° C and 220 RPM to an $OD_{600}$ of 0.65. Protein expression was induced with 0.5 mM IPTG at 18 ° C and 220 RPM for 18 hours. Cells were harvested by centrifugation and stored at -20 ° C until purification.

***Protein purification***: Cells were resuspended in buffer (20 mM Tris HCl, 500 mM NaCl, and 5 mM imidazole at pH = 7.5) with Pierce Protease Inhibitor Tablets (Thermofisher Scientific) until homogenous. Cells were lysed through sonification followed by centrifugation and filtration. Cell lysate was purified using Ni-NTA affinity column (Cytiva) with running buffer (20 mM Tris HCl, 500 mM NaCl, and 5 mM imidazole at pH = 7.5) and elution buffer (20 mM Tris HCl, 100 mM NaCl, and 300 mM imidazole at pH = 7.5). Further purification was performed using a size-exclusion column (HiLoad Superdex 75pg 26/600, Cytiva).

Protein purities were assessed using SDS-PAGE and concentrations were determined using UV-Vis spectroscopy. All protein purities were assessed at > 95% via ImageJ. Protein was exchanged into storage buffer (50 mM Tris HCl, 5% glycerol, pH = 7.5) and stored at 15 mg/mL at -80 ° C.

***Thermal shift assays*****:** To assess the thermostability of PHD2 variants, thermal shift assays were performed. Assays were carried out in a MyiQ2 rt-PCR detection system (Bio-Rad). Protein unfolding was monitored with SYPRO orange (Thermofisher Scientific) using a default FAM excitation and emission protocol. Each well contained 4 µM protein, 1 mM $(NH_4)_2Fe(SO_4)_2$, 1 mM 2OG, and 5X SYPRO orange in 50 mM Tris (pH = 7.5) with a final volume of 40 µL. The temperature was linearly increased by 0.5 ° C every 30 seconds with fluorescence readings being taken every 30 seconds. Melting temperatures ($T_m$) were determined by taking the negative first derivative of the raw fluorescence data and finding the global minimum value.

***Circular dichroism*****:** Circular dichroism (CD) was used to assess the secondary structure of PHD2 variants. Spectra were obtained on a J-815 CD spectropolarimeter (JASCO) using 1 mm cuvette. Data was acquired with a 50 nm/min scan rate, 2 sec D.I.T., 2 nm bandwidth, and 1 nm data pitch. Data was acquired continuously from 255 nm to 195 nm in triplicate for each variant. The HT voltage was maintained below 700 V for the entirety of each scan. Protein was prepared at 5 µM in 10 mM phosphate buffer (pH = 7.5). Raw data was converted to molar ellipticity to correct for protein concentration differences.

***Anaerobic characterization with UV-Vis spectroscopy:*** All spectra were recorded in an anaerobic glove bag with a Cary60 UV-Vis spectrophotometer at 11 ° C. Anaerobic solutions (50 mM Tris, 5% glycerol (pH = 7.5), 50 mM Tris, 50 mM 2OG, 5% glycerol (pH = 7.5), and 0.01 M NaOH) were prepared by flushing argon gas through the solution for two hours followed by equilibration with the anaerobic glove bag environment overnight prior to the experiment. Dry solids for all other compounds were brought into the anaerobic glove bag the day before experiments were performed, and the solutions were prepared with anaerobic solution. To remove ferric iron from purified PHD2 variants, 0.25 mM PHD2 variants were incubated with 2.5 mM EDTA for one hour in the glove bag at 4 ° C. After incubation, the protein was buffer-exchanged four times with anaerobic 50 mM Tris, 5% glycerol (pH = 7.5) in 0.5 mL Amicon Ultra 10 kDa MWCO centricons by centrifugation at 13,500 RPM for 15 minutes at 4 ° C to remove iron-bound EDTA. Solutions of 0.25 mM PHD2 were prepared with 0.2 mM $(NH_4)_2Fe(SO_4)_2$ making the effective PHD2 concentration 0.2 mM.

2OG was added up to 0.25 mM and incubated for 10 minutes to ensure binding equilibrium was established before acquiring spectra. Multiple spectra were taken to ensure the complex was stable.[42] The HIF peptide mimic was also added up to 0.25 mM and spectra were acquired in the same manner. NO solutions were prepared by dissolving proli-NONOate (NO-generating molecule) in 0.01 M NaOH. The solution was added to the protein so that NO concentrations reached ~1 mM upon NO generation. Spectra were acquired as previously stated.

***Stopped-flow UV-Vis spectroscopy:*** All stopped-flow kinetic time traces were acquired on an Applied Photophysics SX-20 system in an anaerobic glove bag. PHD2 variants were prepared in the same manner as previously stated. The final prepared PHD2 solutions were 0.25 mM PHD2, 0.2 mM Fe(II), 0.25 mM 2OG, and 0.25 mM HIF-1α peptide in 50 mM Tris, 5% glycerol (pH = 7.5). Varying concentrated NO solutions were prepared by dissolving varying amounts of DEA-NONOate (NO-generating molecule) in 50 mM Tris, 5% glycerol (pH = 7.5). The solution was aspirated into a Hamilton gas-tight syringe and incubated at 37 $^{\circ}$C for 1 hour to generate NO. NO concentrations generated within the syringe were confirmed using a PreSens Flow-Through Cell PSt3 prior to loading onto the stopped-flow system. This allowed for accurate determination of NO concentrations. Protein solutions and NO solutions were mixed in equal volumes at 11 $^{\circ}$C in the stopped-flow. Kinetic traces were acquired for 25 seconds. Acquired data was fit to a first-order exponential function using Origin 2019 (OriginLab).

***Hydroxylation activity screen*:** Initial hydroxylation activity of PHD2 variants was assessed by monitoring hydroxylation activity at atmospheric oxygen levels. Reactions were carried out at 2 µM PHD variant, 50 µM $(NH_4)_2Fe(SO_4)_2$, 100 µM HIF-1α peptide mimic, 300 µM 2OG, and 4 mM sodium ascorbate in 50 mM Tris buffer (pH = 7.5) at 20 $^{\circ}$C in a 1.7 mL reaction tube. MALDI-TOF mass spectrometry was used to monitor the conversion of the native peptide [(M + $Na^+$), 2276 m/z calculated, 2276 m/z observed] to the hydroxylated peptide [(M + O + $Na^+$), 2292 m/z calculated, 2292 m/z observed]. Reactions were quenched at different time points by spotting 1 µL of reaction mix directly on to the MALDI target plate with 1 µL of saturated 4-α-cyanohydroxycinnamic acid dissolved in 50% acetonitrile and 0.1% trifluoroacetic acid. Spots were analyzed using a positive reflectron method in a Daltonics Autoflex Speed MALDI-TOF mass spectrometer (Bruker). Percent conversion was calculated by comparing the relative intensities of the native and hydroxylated peptide species.

***Steady-state kinetics assays*:** Steady-state constants for PHD2 variants were obtained by monitoring hydroxylation activity at saturating concentrations of 10 µM $(NH_4)_2Fe(SO_4)_2$, 100 µM HIF-1α peptide mimic, 300 µM 2OG, and 2 mM sodium ascorbate in 50 mM Tris (pH = 7.5) at 25 °C. 2 µM of PHD variant was used for these reactions. Reaction was performed in an Oxytherm instrument (Hansatech) with a Clark electrode to monitor oxygen concentration. Concentration of dissolved oxygen was controlled by mixing varying ratios of $N_2$ purged buffer and $O_2$ purged buffer and equilibrating in the Oxytherm reaction vessel. MALDI-TOF mass spectrometry was used to monitor the conversion of the native peptide [(M + $Na^+$), 2276 m/z calculated, 2276 m/z observed] to the hydroxylated peptide [(M + O + $Na^+$), 2292 m/z calculated, 2292 m/z observed]. Reactions were quenched at different time points by spotting 1 µL of reaction mix directly on to the MALDI target plate with 1 µL of saturated 4-α-cyanohydroxycinnamic acid dissolved in 50% acetonitrile and 0.1% trifluoroacetic acid. Samples were analyzed using a positive reflectron method in a Daltonics Autoflex Speed MALDI-TOF mass spectrometer (Bruker).[53] Percent conversion was calculated by comparing the relative intensities of the native and hydroxylated peptide species. Methionine oxidation (~6%) was subtracted, and the data were fitted with the Michaelis-Menten equation.

***Mammalian cell culture and plasmid transfection*:** HEK-293T were cultured in Dulbecco's Modified Eagle's medium (DMEM) (11965092, Gibco) supplemented with 10% Fetal Bovine Serum (F0926, Sigma) and 1% penicillin-streptomycin (Corning). The cells were maintained in a 37 °C incubator supplied with 5% $CO_2$. For plasmid transient transfections, cells were seeded 24 hours before transfection at a density of $1 \times 10^5$ cells/ml, then transfected with 0.2 μg plasmid for the next 24 hours. For hypoxia treatment, 24 hours after transfection, cells were cultured in a hypoxia chamber (BioSperix) in the regular 37 °C incubator supplied with 1% $O_2$/5% $CO_2$/94% $N_2$ for 24 hours. Cell lines were regularly tested for mycoplasma (G238, abm) and found to be negative.

***Cell lysis and Western blotting*:** Cells were harvested by washing with cold PBS and lysed in cell lysis buffer (150 mM NaCl, 0.5% NP-40, 50 mM Tris-HCL, 10% glycerol, 1% SDS, pH 7.5, supplemented with 1x protease inhibitor cocktail (Roche)) on ice. The extracted proteins were separated in homemade SDS-PAGE gel and transferred onto PVDF membrane. Blocking was done with 5% skim milk (BD) in TBST (TBS+0.1% Tween-20).

After blocking, the membrane was incubated with primary antibody overnight at 4°C and washed with TBST for 3 times, then incubated with HRP-linked secondary antibody (Cell Signaling Technology, #7074 and #7076) for 1 hour and washed with TBST. The signal was developed with Luminata Crescendo Western HRP Substrate (WBLUR0500, Millipore) and captured on X-ray film. Primary antibodies used in the current study included Anti-HA (660002, Biolegend), HIF-1α (SAB2702132, Millipore), α-Tubulin (T6199, Sigma). Protein bands were quantified using ImageJ.

***Two-step quantitative reverse transcriptase polymerase chain reaction (RT-qPCR):*** RNA was prepared and isolated from transfected HEK-293T cells using Total RNA Miniprep Kit (New England Biolabs) according to the manufacturers protocol. RNA was reverse transcribed to cDNA using SuperScript IV VILO master mix (ThermoFisher Scientific) according to the manufacturers protocol. qPCR was then performed using Luna Universal qPCR master mix (New England Biolabs) on a MyiQ2 rt-PCR detection system with iQ5 software (Bio-Rad). β-actin (ACTB) was used as the normalizing housekeeping gene. Fold changes were determined using the $\Delta\Delta C_t$ method. All primers were validated, and the sequences were as follows: ACTB (forward – GCTGTGCTACGTCGCCCTG, reverse – GGAGGAGCTGGAAGCAGCC), ADM (forward – TTGGCAGATCACTCTCTTAG, reverse – TTCCACTTCTTTCGAAACTC), HK2 (forward – CCCCTGCCACCAGACTAAACTA, reverse – CAAAGTCCCCTCTCCTCTGGAT), LDHA (forward – CACCATGATTAAGGGTCTTTAC, reverse – AGGTCTGAGATTCCATTCTG). Genes were analyzed through three independent experiments with three technical replicates each experiment.

**Table S1. $O_2$ kinetic constants of oxygen sensing non-heme iron hydroxylases**

| Protein | $K_M(O_2)$ (μM) | $k_{cat}$ ($s^{-1}$) | $k_{cat}/K_M(O_2)$ ($M^{-1}$ $s^{-1}$) | Species | Method | Reference |
|---|---|---|---|---|---|---|
| AspH | 426 ± 73 | 0.23 ± 0.04 | 540 ± 130 | *H. sapien* | SPE MS | 6 |
| FIH | 110 ± 30 | 0.56 ± 0.04 | 5090 ± 1400 | *H. sapien* | MALDI-TOF MS | 7 |
| TgPhyA | 2.7 ± 0.54 | NA | NA | *T. gondii* | SPE MS | 8 |
| DdPhyA | > 810 | NA | NA | *D. discoideum* | SPE MS | 8 |
| PPHD | 47.5 ± 9.2 | 0.018 ± 0.006 | 380 ± 150 | *P. putida* | MALDI-TOF MS | 10 |
| PHD | > 400 | NA | NA | *T. adhaerans* | MALDI-TOF MS | 11 |
| PHD2 | 460 ± 30<br>229 ± 60<br>530 ± 90<br>415 ± 36 | 0.060 ± 0.006<br>0.040 ± 0.004<br>0.13 ± 0.01<br>0.037 ± 0.002 | 130 ± 16<br>175 ± 49<br>245 ± 46<br>89 ± 9 | *H. sapien* | MALDI-TOF MS | 7<br>8<br>38<br>This work |

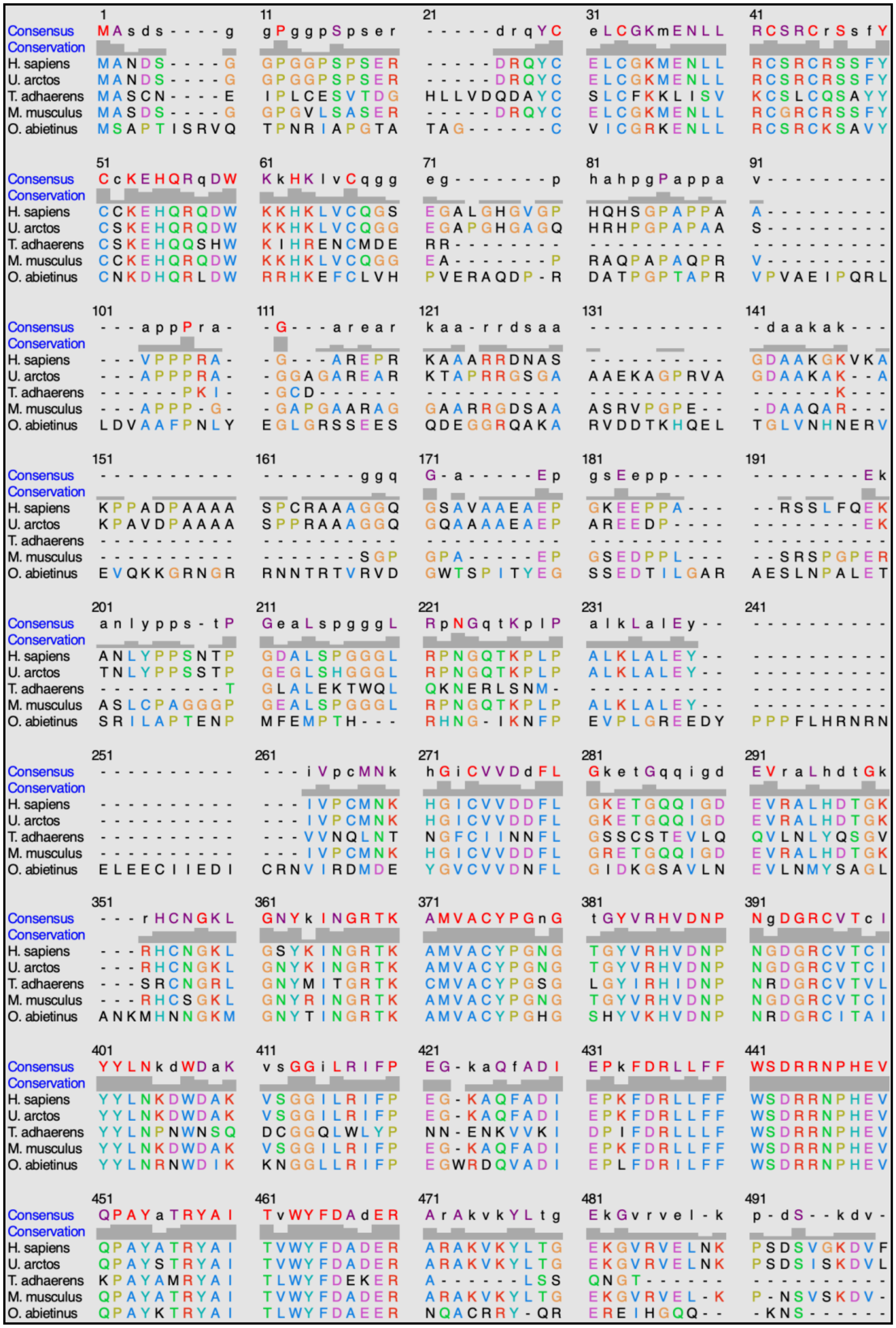

**Figure S1.** Amino acid alignment of *H. sapiens* PHD2 with other eukaryotic PHD homologs (*U. arctos, T. adhaerens, M. musculus, O. abietinus*). Conservation of each residue positively correlates with the height of the gray bar above each residue. The N-terminus zinc finger domain corresponds to residues 21-71 and the C-terminus hydroxylase domain corresponds to residues 261-500.

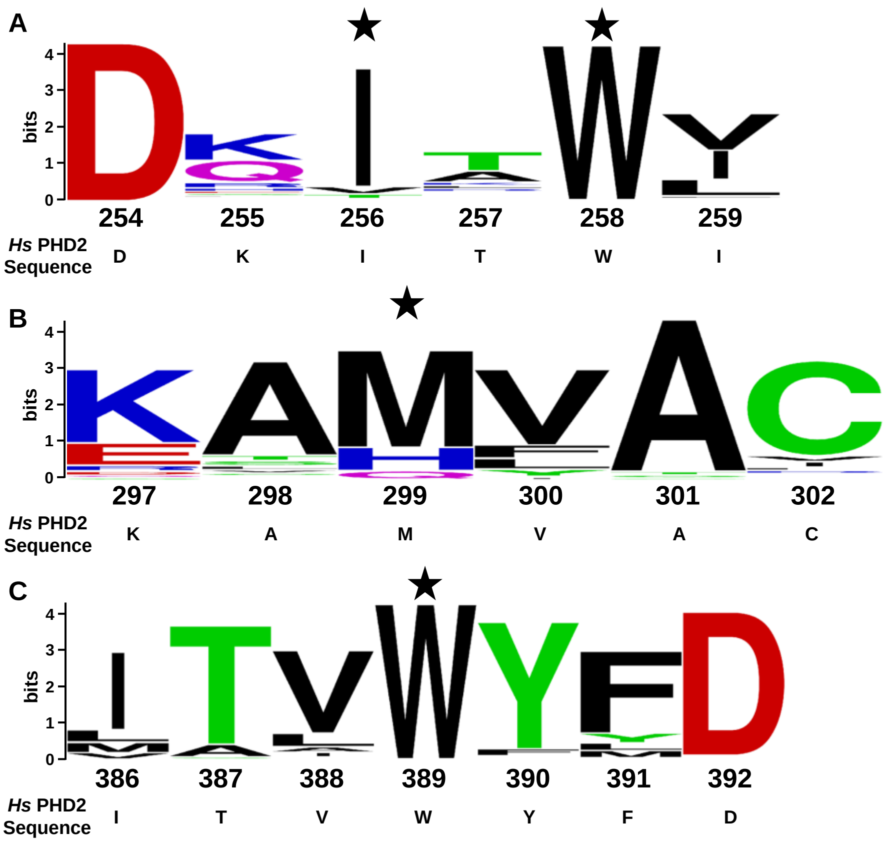


**Figure S2.** Logo plots representing the conservation and frequency of tunnel forming amino acids in homologous PHDs. The height of each column represents the conservation of amino acids at each position, whereas the height of each letter represents the frequency of a specific amino acid at each position. Starred residues indicate which amino acids were selected for mutagenesis. The primary amino acid sequence and numbering of *Homo sapiens* PHD2 can be found below each logo plot.

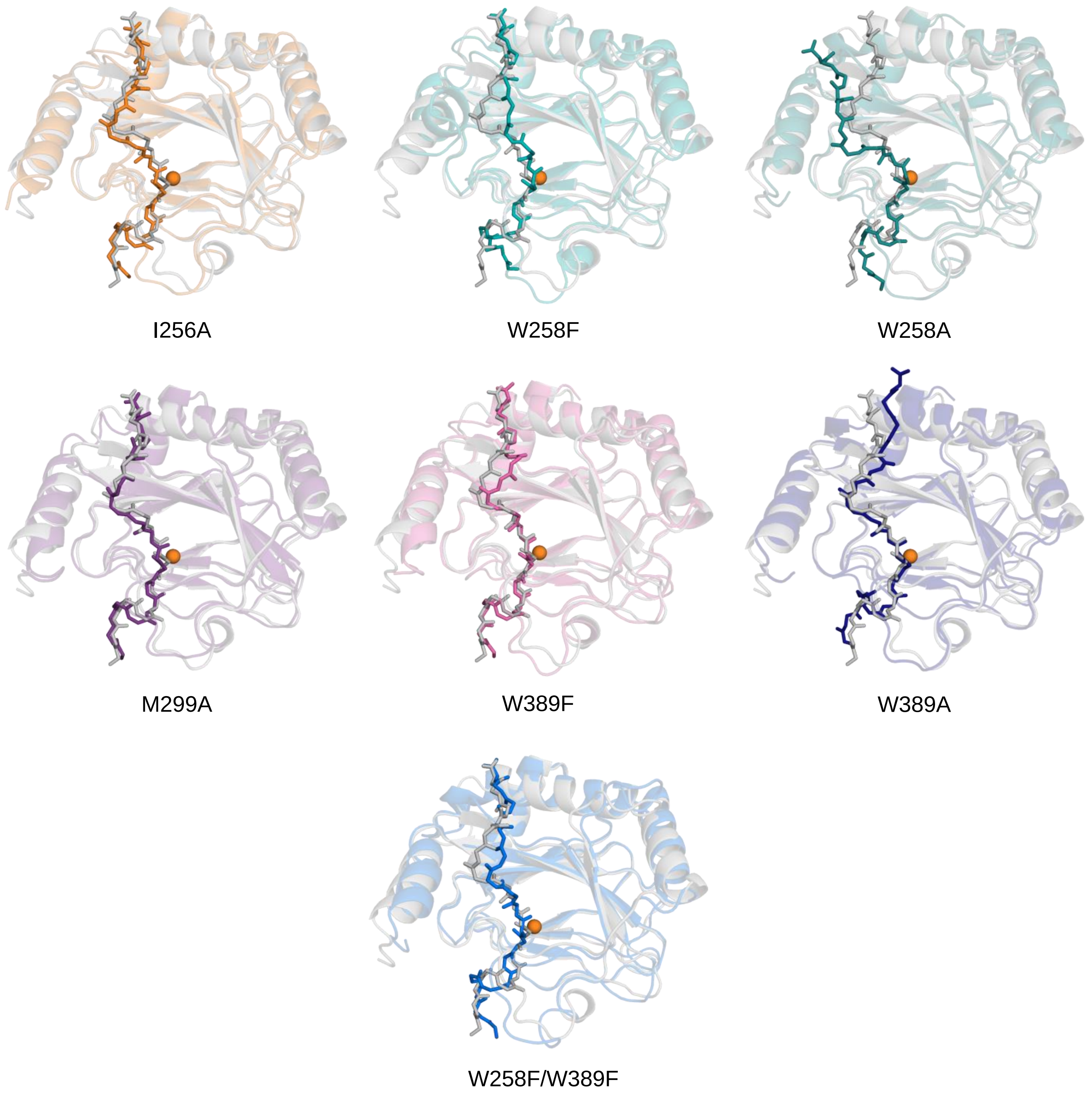


**Figure S3.** Representative conformations of PHD2 and HIF-1α peptide variants from MD simulations. HIF-1α peptide is shown in opaque whereas PHD2 is shown transparent. PHD2 variants (colored) are overlaid with WT PHD2 (gray) to show comparison between variant and WT structures.

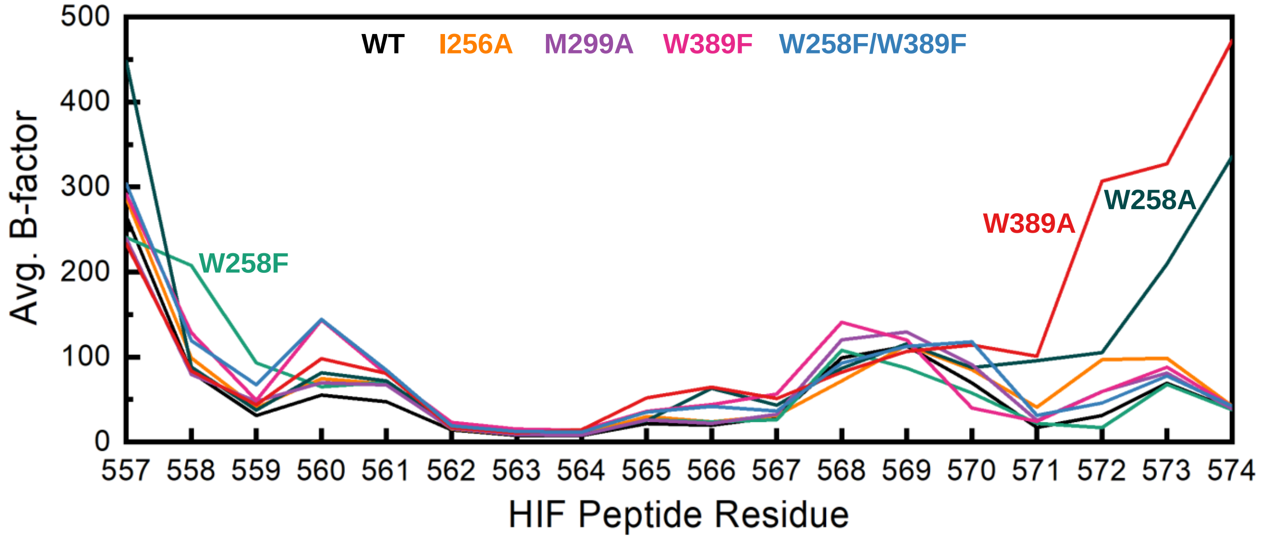


**Figure S4.** B-factor of HIF-1α peptide residues from MD simulations. Higher B-factor values are associated with increased fluctuation.

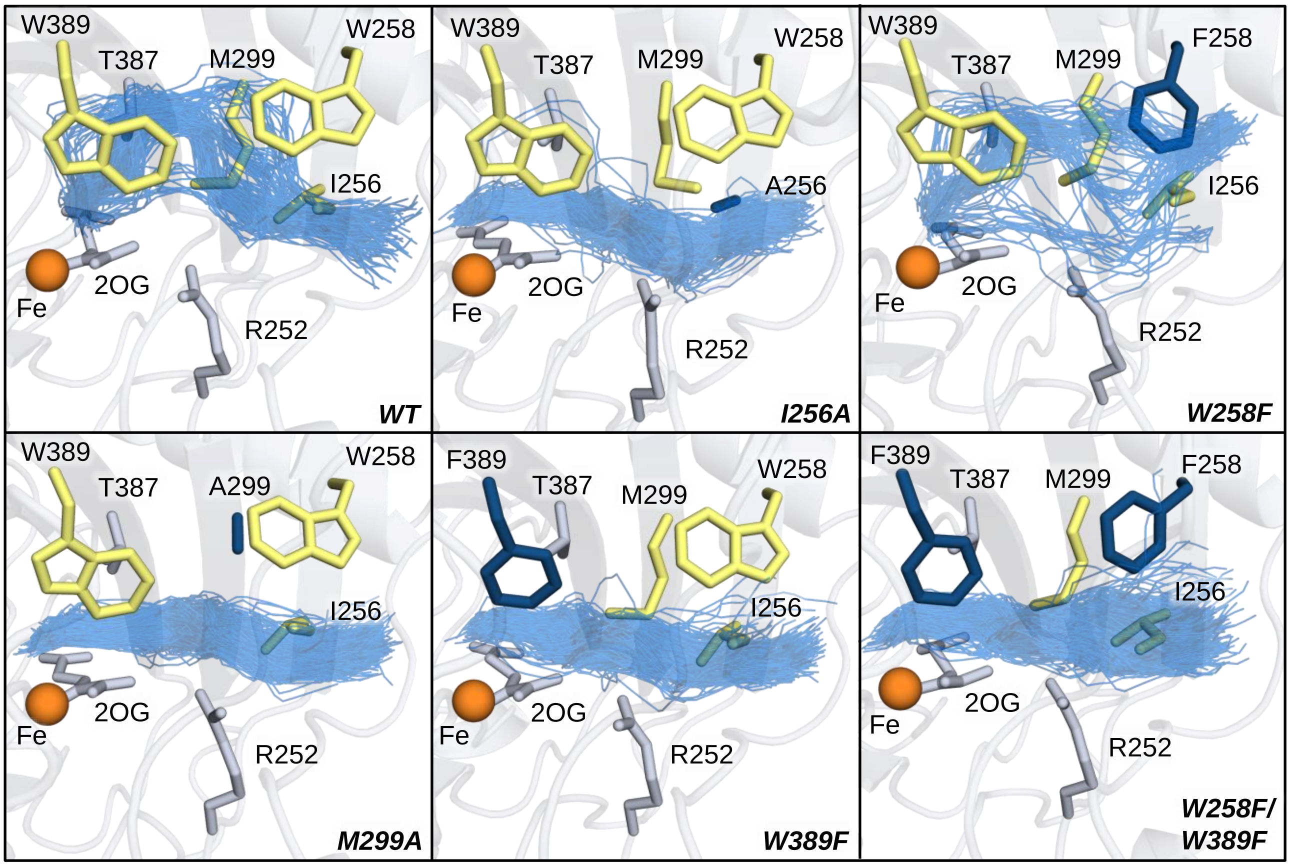


**Figure S5.** Traces of CAVER computed tunnel paths (blue lines) from MD simulations. Each line represents the tunnel that was found in a specific frame over the course of the simulation. WT tunnel forming residues that were targets of our rational design studies are shown in yellow in each panel. Residues that were mutated are shown in dark blue. Residues that potentially constricted the tunnel but were not targets of mutagenesis are shown in gray.

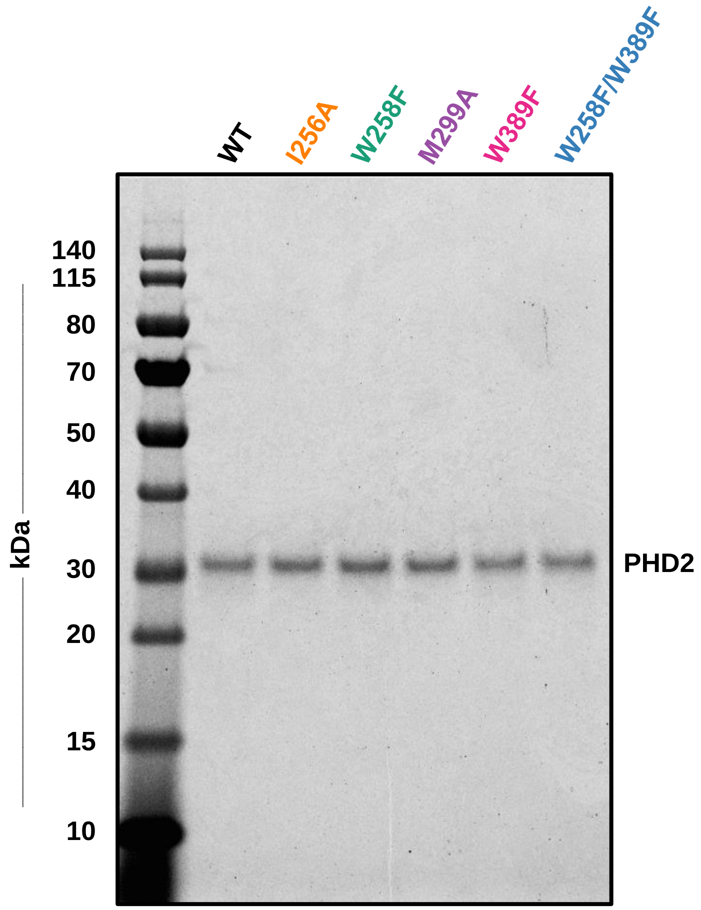


**Figure S6.** SDS-PAGE gel for purified PHD2 variants after IMAC and SEC. All variants have molecular weight of ~29.9 kDa, and all samples have >95% purity as quantified by ImageJ.

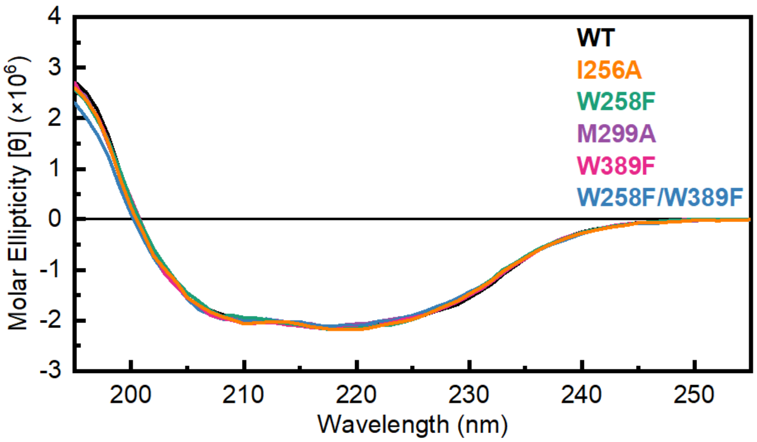


**Figure S7.** Circular dichroism spectra of PHD2 variants. Mutated PHDs show no significant differences in secondary structure compared to wild-type PHD. The spectra are the average of three independent scans, and they were converted to molar ellipticity to correct for variation in protein concentration.

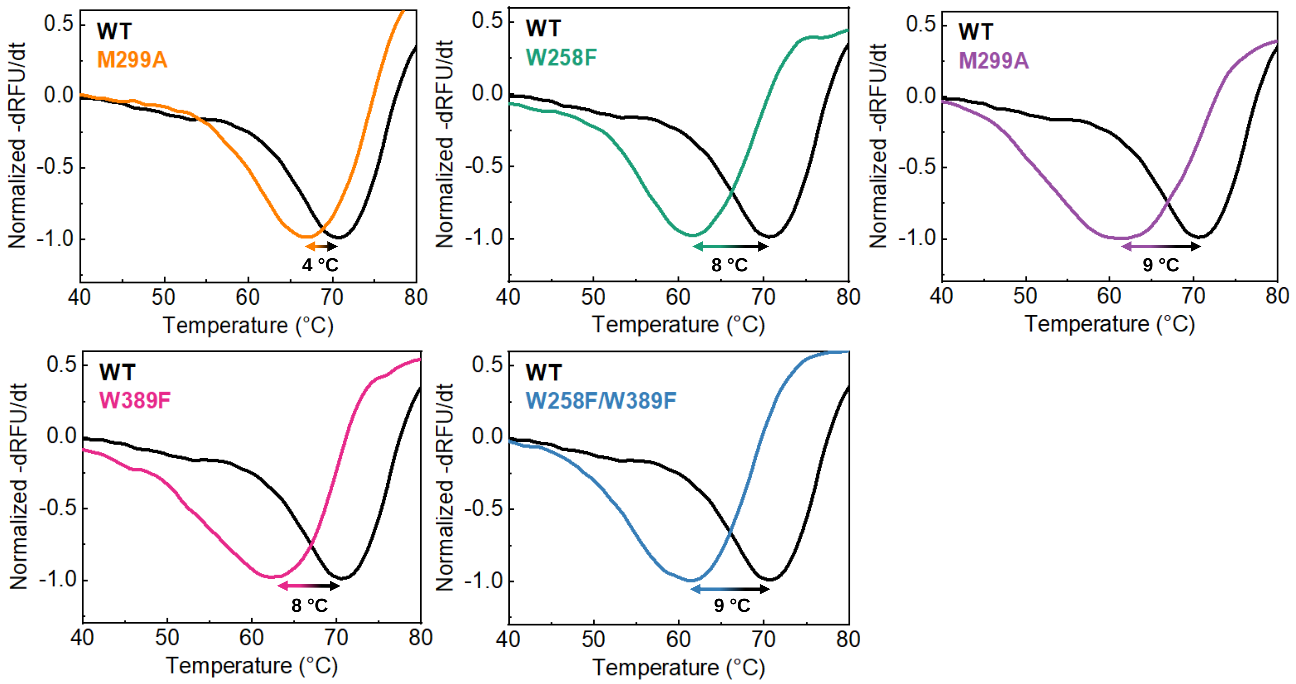


**Figure S8.** Thermal shift assays of PHD2 variants. All variants exhibit a lower melting temperature ($T_m$) compared to WT ($T_m$ = 70.5 °C). The $T_m$ difference between WT and variants can be found in each panel. Each curve is the average of three independent samples, and the data is smoothed every 5 data points using a Savitzky-Golay filter.

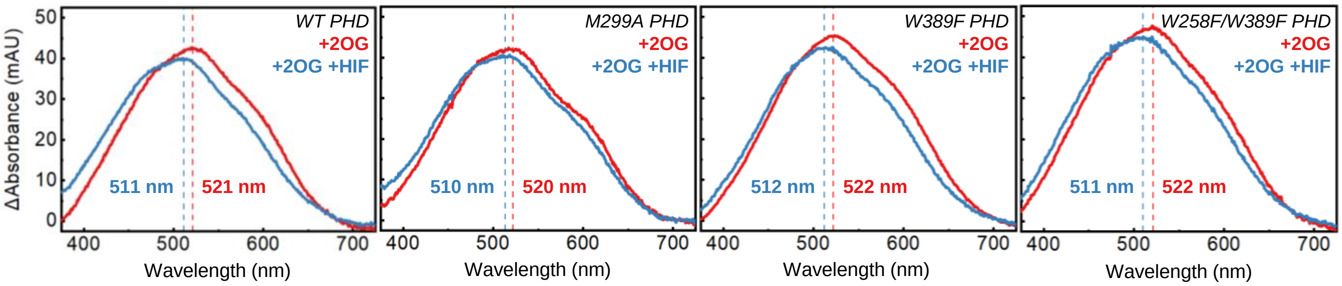


**Figure S9.** Difference UV-Vis spectra of PHD2 variant active-site assembly and MLCT perturbation**.** 2OG-bound PHD2 exhibits a broad MLCT centered around 520 nm (red). Binding of the HIF-1α peptide forces the loss of an aqua ligand resulting in a blue-shift of the MLCT (blue).

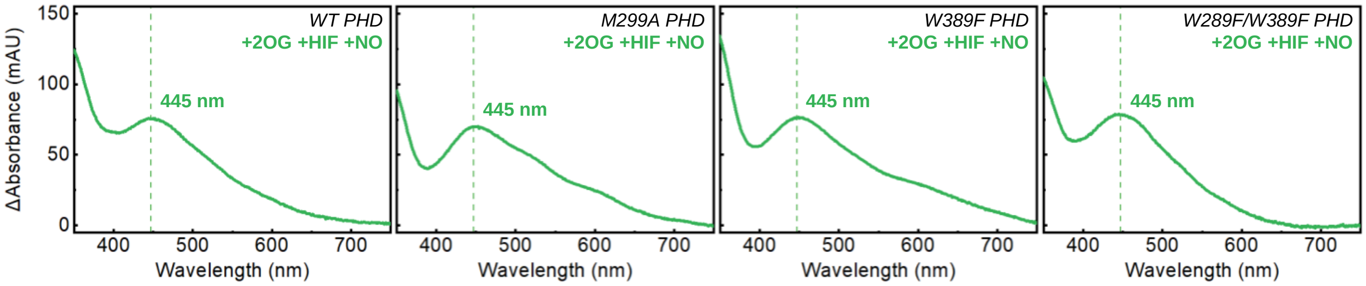


**Figure S10.** Difference UV-Vis spectra of iron nitrosyl formation in PHD2 variants. After assembling the active site with 2OG and the HIF-1α peptide, exposing the PHD2 variants to NO forms an iron nitrosyl complex which exhibits a distinct UV-Vis signature at 445 nm.

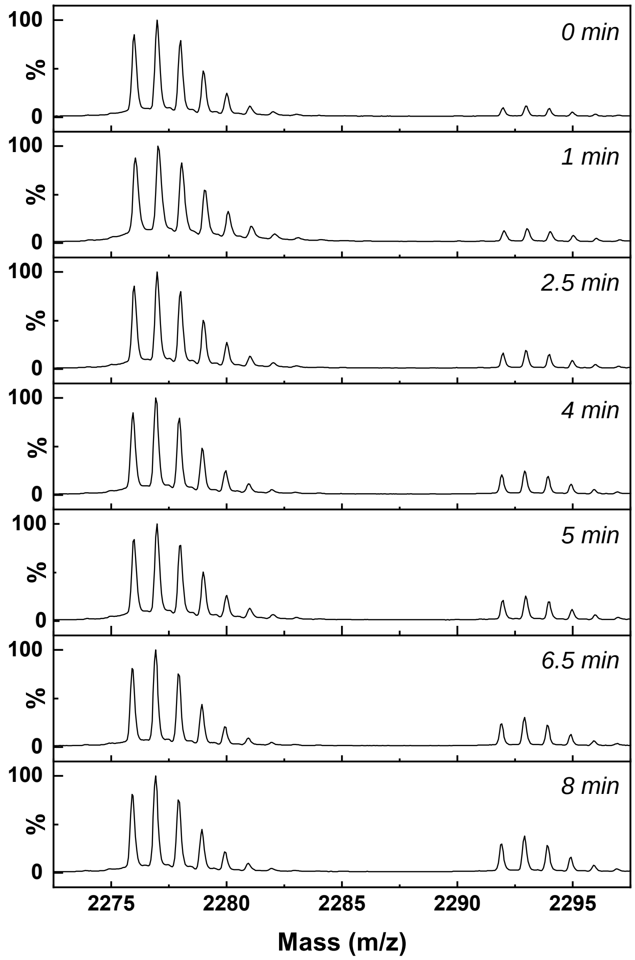


**Figure S11.** Monitoring HIF-1α peptide hydroxylation over time using MALDI-TOF MS. HIF-1α peptide [(M + $Na^+$), 2276 m/z] is hydroxylated [(M + O + $Na^+$), 2292 m/z] indicated by the increasing intensity of the hydroxylated HIF-1α peptide as time increases. Each grouping of peaks are isotopologs of their respective species.

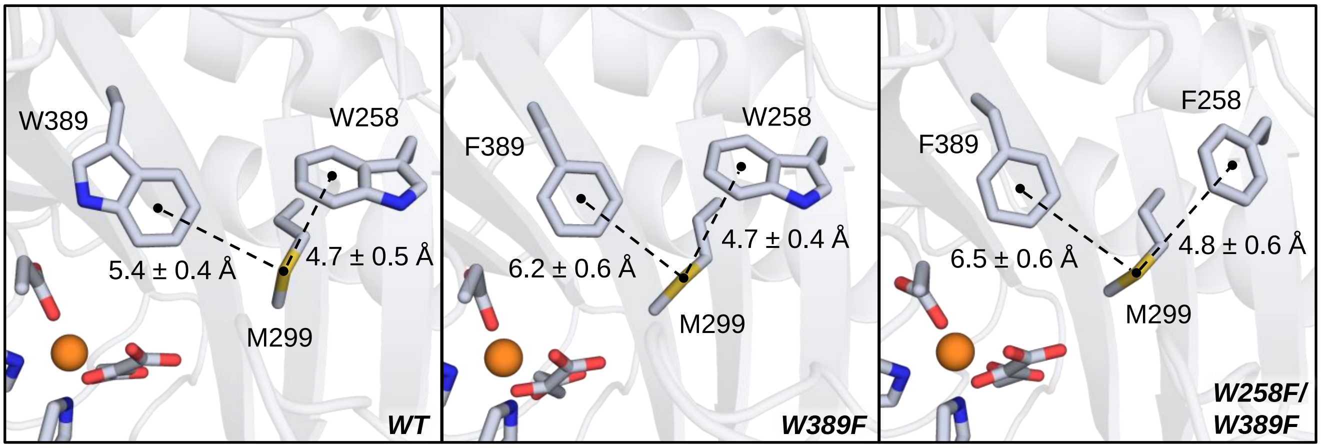


**Figure S12.** Aromatic-methionine bridge motif observed in PHD2 variants. Distances are averages calculated from MD simulations. Mutations induced in the W389F and W258F/W389F variants maintain the amino acid properties and distances needed for these interactions to form.

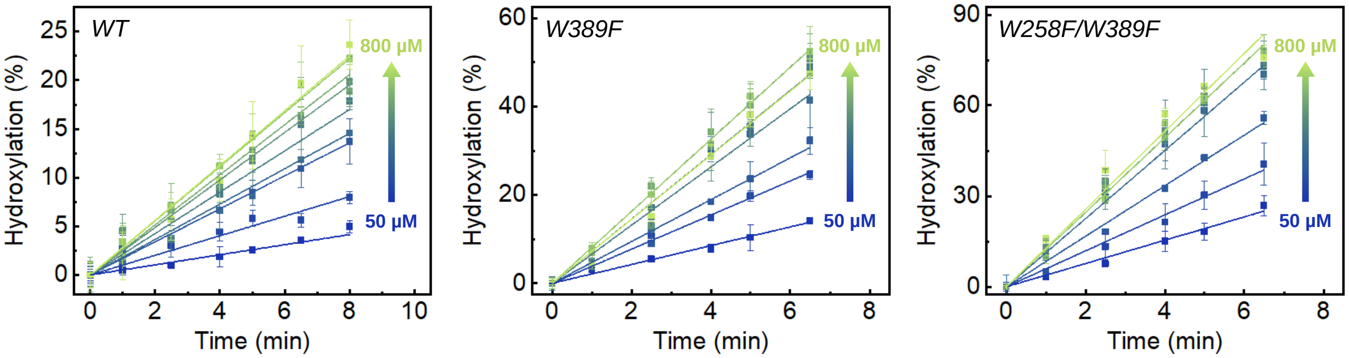


**Figure S13.** Hydroxylation activity time traces of PHD2 variants at varying $O_2$ concentrations (50-800 µM $O_2$). Hydroxylation activity increases as a function of increasing oxygen concentration where low oxygen concentration trials are colored blue and high oxygen concentration trials are colored light green.

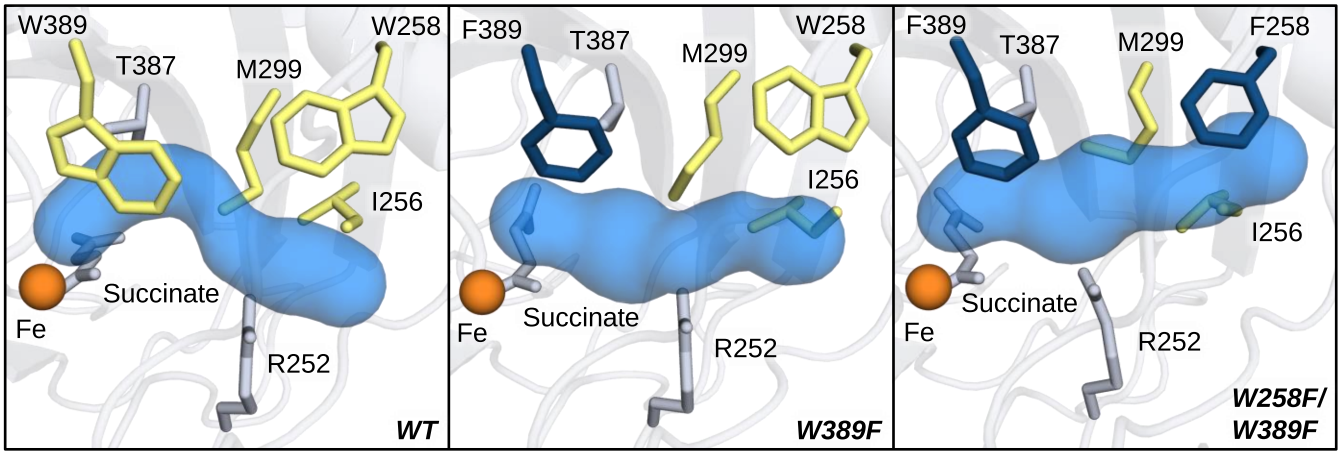


**Figure S14.** Representative structures from MD simulations of ferryl-intermediate PHD2 variants. The CAVER computed $CO_2$ tunnels are shown in blue. WT tunnel forming residues that were targets of mutagenesis are shown in yellow in each panel. Residues that were mutated are shown in dark blue. Residues that potentially constricted the tunnel but were not targets of mutagenesis are shown in gray.

**Table S2. CAVER statistics of ferryl-intermediate PHD2 tunnel variants for $CO_2$ egress from iron center**

| Variant | % of frames with primary tunnel (Avg.) ± SD (%) | Avg. BR ± AD (Å) | Avg. Length ± AD (Å) | Avg. Curvature ± AD (Å) |
|---|---|---|---|---|
| WT | 51.3 ± 5.5 | 1.03 ± 0.11 | 15.6 ± 2.2 | 1.33 ± 0.14 |
| W389F | 67.3 ± 36.4 | 1.06 ± 0.13 | 15.5 ± 2.2 | 1.25 ± 0.13 |
| W258F/W389F | 88.0 ± 12.1 | 1.17 ± 0.15 | 15.1 ± 1.9 | 1.22 ± 0.10 |

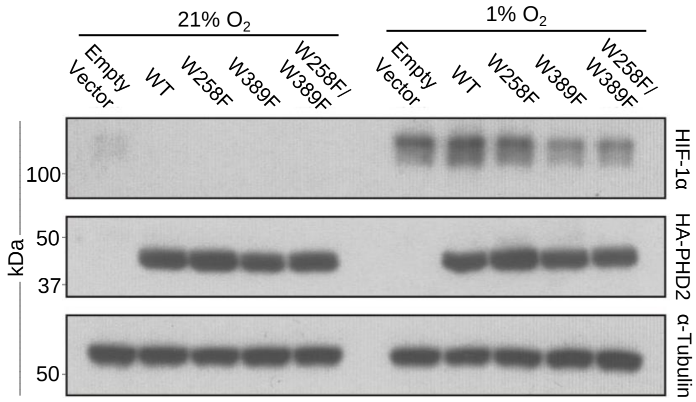


**Figure S15.** Western blot from hypoxia signaling experiments. HEK-293T cells transfected with PHD variants were incubated under normoxia (21% $O_2$, left of blot) and extreme hypoxia (1% $O_2$, right of blot). Blots were stained with primary antibodies, followed by an HRP-linked secondary antibody. Images were captured on X-ray film. Three independent experiments were completed for quantification.

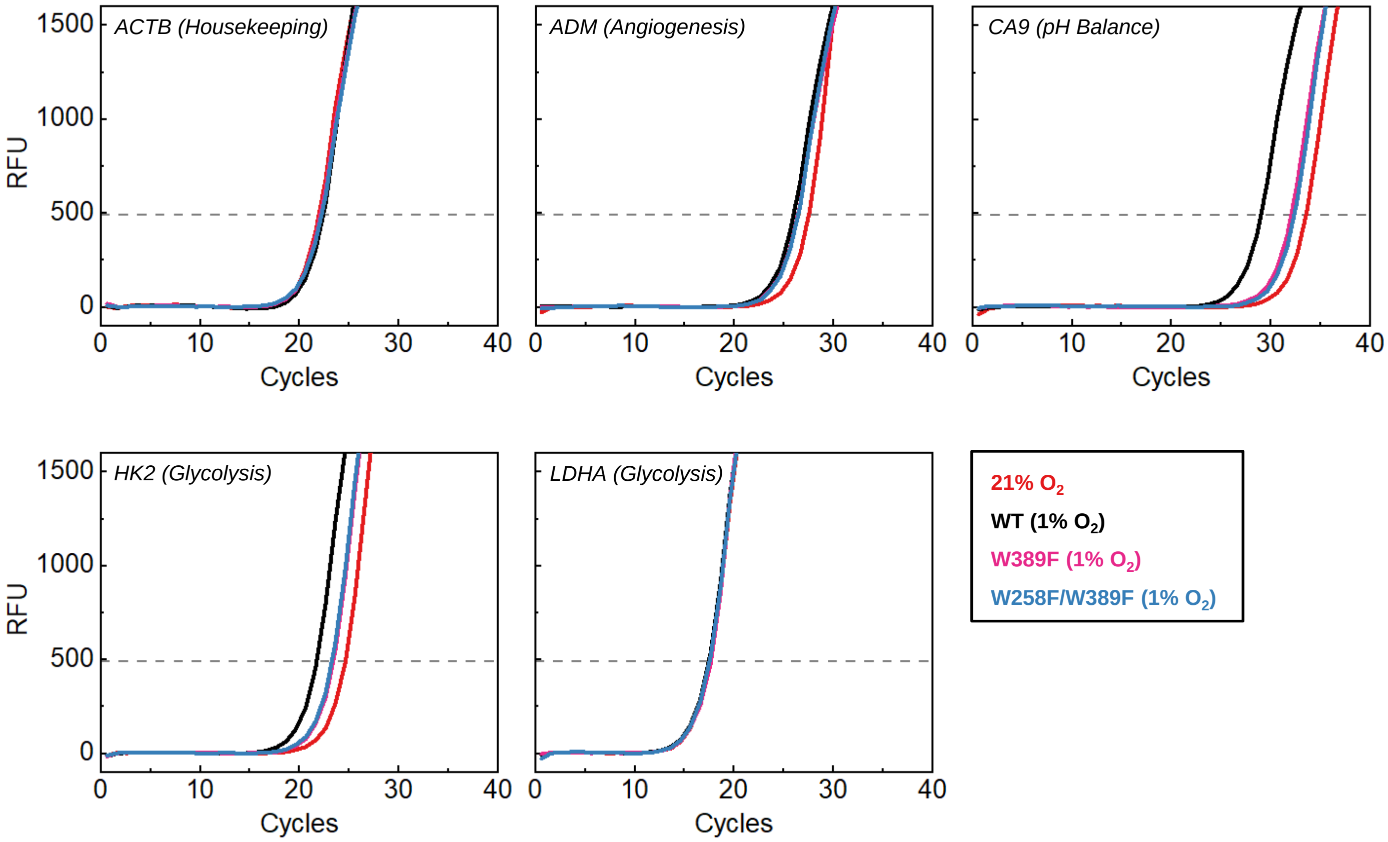


**Figure S16.** Average fluorescence amplification curves from RT-qPCR experiments. Each curve is the average of nine data sets (three independent experiments with three technical replicates each). $C_t$ threshold (gray dash line) was determined using the iQ5 software.
